# Integrative proteomics and bioinformatic prediction enable a high-confidence apicoplast proteome in malaria parasites

**DOI:** 10.1101/265967

**Authors:** Michael J. Boucher, Sreejoyee Ghosh, Lichao Zhang, Avantika Lal, Se Won Jang, An Ju, Shuying Zhang, Xinzi Wang, Stuart A. Ralph, James Zou, Joshua E. Elias, Ellen Yeh

**Affiliations:** Department of Microbiology and Immunology, Stanford University School of Medicine, Stanford, CA 94305, United States of America; Department of Biochemistry, Stanford University School of Medicine, Stanford, CA 94305, United States of America; Department of Chemical and Systems Biology, Stanford University School of Medicine, Stanford, CA 94305, United States of America; Department of Pathology, Stanford University School of Medicine, Stanford, CA 94305, United States of America; Department of Computer Science, Stanford University, Stanford, CA 94305, United States of America; Department of Bioengineering, Stanford University, Stanford, CA 94305, United States of America; Department of Biochemistry and Molecular Biology, Bio21 Molecular Science and Biotechnology Institute, The University of Melbourne, Parkville, Vic 3010, Australia; Department of Biomedical Data Science, Stanford University School of Medicine, Stanford, CA 94305, United States of America; Chan Zuckerberg Biohub, San Francisco, CA 94158, United States of America

**Keywords:** malaria, apicoplast, BioID, proteome, neural network

## Abstract

Malaria parasites *(Plasmodium* spp.) and related apicomplexan pathogens contain a non-photosynthetic plastid called the apicoplast. Derived from an unusual secondary eukaryote-eukaryote endosymbiosis, the apicoplast is a fascinating organelle whose function and biogenesis rely on a complex amalgamation of bacterial and algal pathways. Because these pathways are distinct from the human host, the apicoplast is an excellent source of novel antimalarial targets. Despite its biomedical importance and evolutionary significance, the absence of a reliable apicoplast proteome has limited most studies to the handful of pathways identified by homology to bacteria or primary chloroplasts, precluding our ability to study the most novel apicoplast pathways. Here we combine proximity biotinylation-based proteomics (BioID) and a new machine learning algorithm to generate a high-confidence apicoplast proteome consisting of 346 proteins. Critically, the high accuracy of this proteome significantly outperforms previous prediction-based methods and extends beyond other BioID studies of unique parasite compartments. Half of identified proteins have unknown function, and 77% are predicted to be important for normal blood-stage growth. We validate the apicoplast localization of a subset of novel proteins and show that an ATP-binding cassette protein ABCF1 is essential for blood-stage survival and plays a previously unknown role in apicoplast biogenesis. These findings indicate critical organellar functions for newly discovered apicoplast proteins. The apicoplast proteome will be an important resource for elucidating unique pathways derived from secondary endosymbiosis and prioritizing antimalarial drug targets.

## Introduction

Identification of new antimalarial drug targets is urgently needed to address emerging resistance to all currently available therapies. However, nearly half of the *Plasmodium falciparum* genome encodes conserved proteins of unknown function [1], obscuring critical pathways required for malaria pathogenesis. The apicoplast is an essential, non-photosynthetic plastid found in *Plasmodium* spp. and related apicomplexan pathogens [2, 3]. This unusual organelle is an enriched source of both novel cellular pathways and parasite-specific drug targets [4]. It was acquired by secondary (i.e., eukaryote-eukaryote) endosymbiosis and has evolutionarily diverged from the primary endosymbiotic organelles found in model organisms. While some aspects of apicoplast biology are shared with bacteria, mitochondria, and primary chloroplasts, many are unique to the secondary plastid in this parasite lineage. For example, novel translocons import apicoplast proteins through specialized membranes derived from secondary endosymbiosis [5–8], while the parasite’s pared-down metabolism necessitates export of key metabolites from the apicoplast using as-yet unidentified small molecule transporters [9, 10].

These novel cellular pathways, which are also distinct from human host cells, can be exploited for antimalarial drug discovery. Indeed, antimalarials that target apicoplast pathways are currently in use as prophylactics or partner drugs (doxycycline, clindamycin) or have been tested in clinical trials (fosmidomycin) [11–15]. However, known apicoplast drug targets have been limited to the handful of pathways identified by homology to plastid-localized pathways in model organisms. Meanwhile the number of druggable apicoplast targets, including those in unique secondary plastid pathways, is likely more extensive [16].

A major hurdle to identifying novel, parasite-specific pathways and prioritizing new apicoplast targets is the lack of a well-defined organellar proteome. So far, the apicoplast has not been isolated in sufficient yield or purity for traditional organellar proteomics. Instead, large-scale, unbiased identification of apicoplast proteins has relied on bioinformatic prediction of apicoplast targeting sequences [17–19]. These prediction algorithms identify hundreds of putative apicoplast proteins but contain numerous false positives. Confirmation of these low-confidence candidate apicoplast proteins is slow due to the genetic intractability of *P. falciparum* parasites. Unbiased identification of apicoplast proteins in an accurate and high-throughput manner would significantly enhance our ability to study novel apicoplast pathways and validate new antimalarial drug targets.

BioID and other cellular proximity labeling methods are attractive techniques for identification of organellar proteins [20, 21]. In BioID, a promiscuous biotin ligase, BirA*, is fused to a bait protein and catalyzes biotinylation of neighbor proteins in intact cells. Proximity labeling methods have been used for unbiased proteomic profiling of subcellular compartments in diverse parasitic protozoa, including *Plasmodium* spp. [22–29]. Here we used BioID to perform large-scale identification of *P. falciparum* apicoplast proteins during asexual blood-stage growth. Extending beyond previous BioID studies of unique parasite compartments, we achieved high positive predictive value of true apicoplast proteins by implementing an endoplasmic reticulum (ER) negative control to remove frequent contaminants expected based on the trafficking route of apicoplast proteins. Furthermore, higher coverage was achieved by using the proteomic dataset to develop an improved neural network prediction algorithm, PlastNN. We now report a high-confidence apicoplast proteome of 346 proteins rich in novel and essential functions.

## Results

### The promiscuous biotin ligase BirA* is functional in the *P. falciparum* apicoplast and endoplasmic reticulum

To target the promiscuous biotin ligase BirA* to the apicoplast, the *N*-terminus of a GFP-BirA* fusion protein was modified with the apicoplast-targeting leader sequence from acyl carrier protein (ACP) (Fig 1A). Since apicoplast proteins transit the parasite ER en route to the apicoplast [30], we also generated a negative control in which GFP-BirA* was targeted to the ER via an *N*-terminal signal peptide and a *C*-terminal ER-retention motif (Fig 1A). Each of these constructs was integrated into an ectopic locus in Dd2^attB^ parasites [31] to generate BioID-Ap and BioID-ER parasites (S1A Fig). Immunofluorescence co-localization and live imaging of these parasites confirmed GFP-BirA* localization to either the apicoplast or the ER, respectively (Fig 1B and S1B Fig).

**Fig 1.**
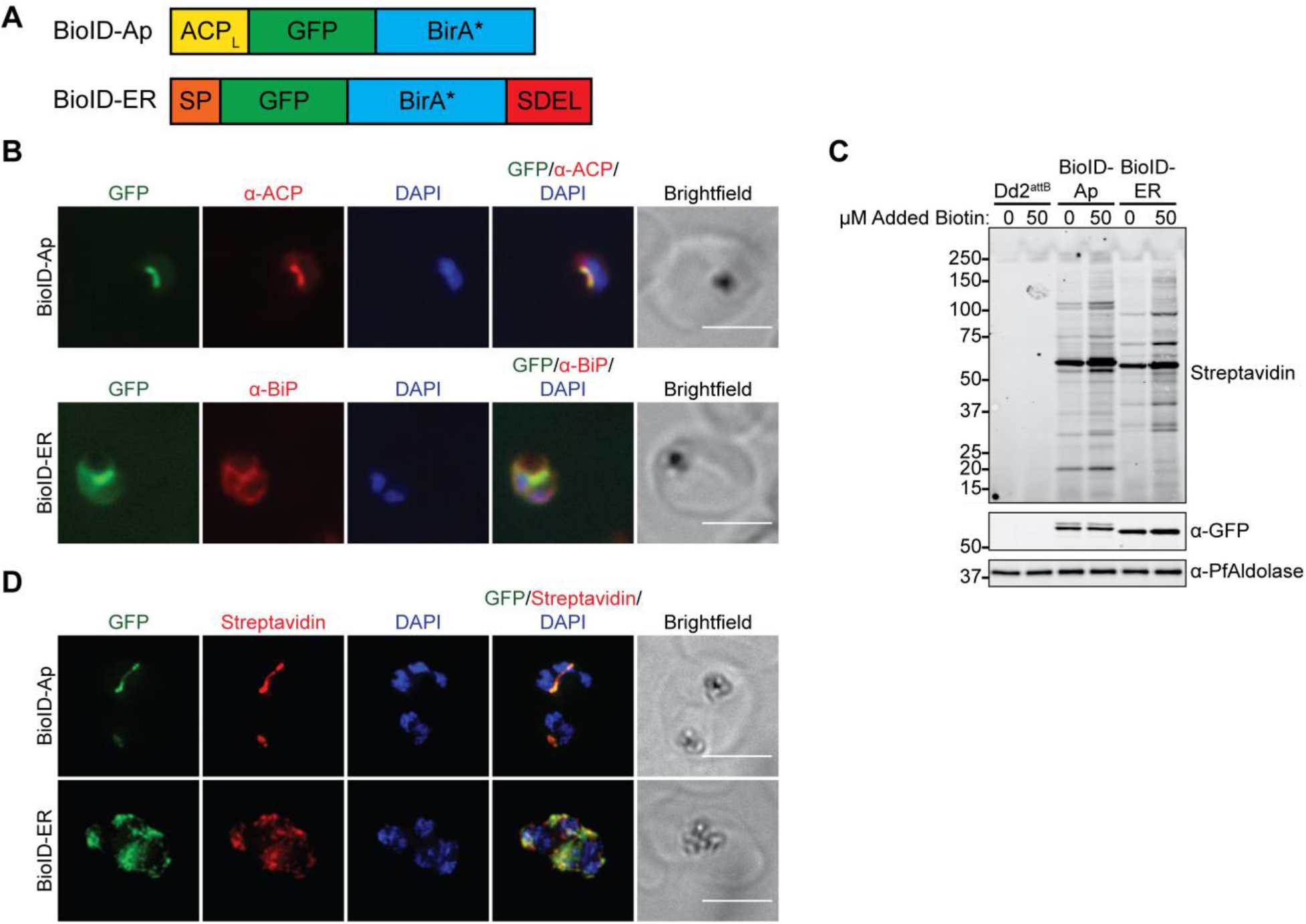
The promiscuous biotin ligase BirA* biotinylates proteins in the *P. falciparum* apicoplast and ER. (A) Schematic (not to scale) of constructs for apicoplast- and ER-targeting of GFP-BirA*. ACP_L_, ACP leader sequence; SP, signal peptide; SDEL, ER-retention motif. (B) Fixed-cell imaging of BioID-Ap and BioID-ER parasites stained with antibodies raised against the apicoplast marker ACP or the ER marker BiP, respectively. Scale bars, 5 μm. (C) Western blot of untreated and biotin-labeled Dd2^attB^, BioID-Ap, and BioID-ER parasites. (D) Fixed-cell imaging of biotinylated proteins in biotin-labeled BioID-Ap and BioID-ER parasites. Scale bars, 5 μm.

To test the functionality of the GFP-BirA* fusions in the apicoplast and ER, we labeled either untransfected Dd2^attB^, BioID-Ap, or BioID-ER parasites with DMSO or 50 μM biotin and assessed biotinylation by western blotting and fixed-cell fluorescent imaging. As has been reported [28], significant labeling of GFP-BirA*-expressing parasites above background was achieved even in the absence of biotin supplementation, suggesting that the 0.8 μM biotin in RPMI growth medium is sufficient for labeling (Fig 1C). Addition of 50 μM biotin further increased protein biotinylation. Fluorescence imaging of biotinylated proteins revealed staining that co-localized with the respective apicoplast-or ER-targeted GFP-BirA* fusion proteins (Fig 1D). These results confirm that GFP-BirA* fusions are active in the *P. falciparum* apicoplast and ER and can be used for compartment-specific biotinylation of proteins.

### Proximity-dependent labeling (BioID) generates an improved apicoplast proteome dataset

For large-scale identification of apicoplast proteins, biotinylated proteins from late-stage BioID-Ap and BioID-ER parasites were purified using streptavidin-conjugated beads and identified by mass spectrometry. A total of 728 unique *P. falciparum* proteins were detected in the apicoplast and/or ER based on presence in at least 2 of 4 biological replicates and at least 2 unique spectral matches in any single mass spectrometry run (Fig 2A and S1 Table). The abundance of each protein in apicoplast and ER samples was calculated by summing the total MS1 area of all matched peptides and normalizing to the total MS1 area of all detected *P. falciparum* peptides within each mass spectrometry run.

**Fig 2.**
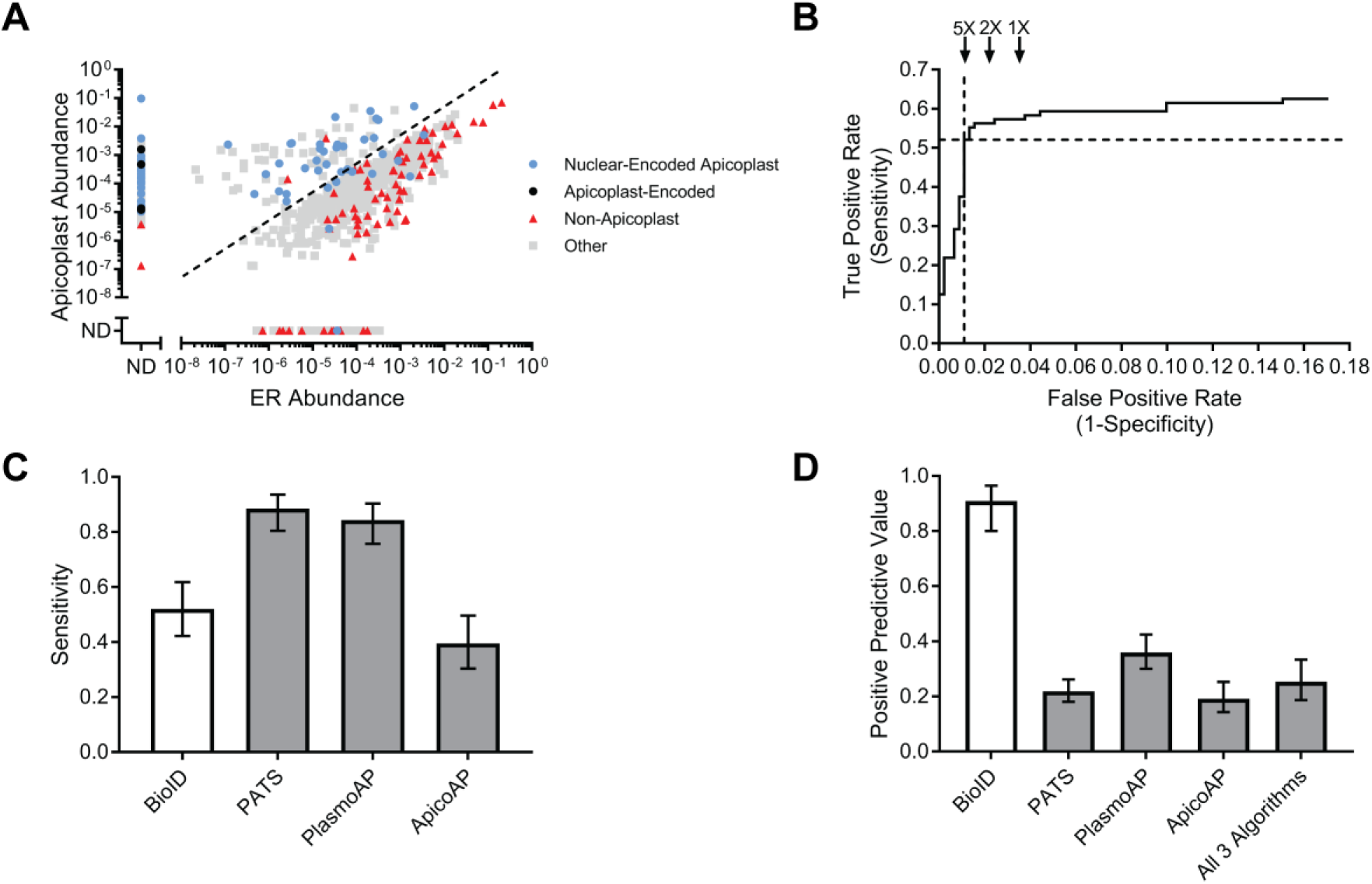
Accurate, unbiased identification of apicoplast proteins using BioID. (A) Abundances of 728 proteins identified by mass spectrometry in BioID-Ap and BioID-ER parasites. Protein abundances were calculated by summing the total MS1 area of all matched peptides for a given protein and normalized by the total summed intensity of all *P. falciparum* peptides matched. Dotted line represents 5-fold apicoplast:ER enrichment. ND, not detected. (B) ROC curve used to identify the apicoplast:ER enrichment that maximized true positives while minimizing false positives. Dotted lines denote the sensitivity and false positive rate of the 5-fold cutoff used. False positive rates for hypothetical 2-fold and 1-fold enrichments are shown for reference. (C) Sensitivities of BioID, PATS, PlasmoAP, and ApicoAP based on identification of 96 known apicoplast proteins. (D) PPV of BioID, PATS, PlasmoAP, ApicoAP, and a dataset consisting of proteins predicted to localize to the apicoplast by all three bioinformatic algorithms. Calculated as the number of true positives divided by the total number of true positives and false positives. Error bars in (C) and (D) represent 95% confidence intervals.

To assess the ability of our dataset to distinguish between true positives and negatives, we generated control lists of 96 known apicoplast and 451 signal peptide-containing non-apicoplast proteins based on published localizations and pathways (S2 Table). Consistent with an enrichment of apicoplast proteins in BioID-Ap samples, we observed a clear separation of known apicoplast and non-apicoplast proteins based on apicoplast:ER abundance ratio (Fig 2A). Using receiver operating characteristic (ROC) curve analysis (Fig 2B), we set a threshold of apicoplast:ER abundance ratio ≥5-fold for inclusion of 187 proteins in the BioID apicoplast proteome, which maximized sensitivity while minimizing false positives (Fig 2A, dotted line; S1 Table). This dataset included 50 of the 96 positive control proteins for a sensitivity of 52% (95% CI: 42-62%). None of the original 451 negative controls were present above the ≥5-fold enrichment threshold, but manual inspection of this list identified 5 likely false positives not present on our initial list (S1 Table) for a positive predictive value (PPV; the estimated fraction of proteins on the list that are true positives) of 91% (95% CI: 80-96%).

To benchmark our dataset against the current standard for large-scale identification of apicoplast proteins, we compared the apicoplast BioID proteome to the predicted apicoplast proteomes from three published bioinformatic algorithms: PATS [17], PlasmoAP [18], and ApicoAP [19] (S3 Table). At 52% sensitivity, apicoplast BioID identified fewer known apicoplast proteins than PATS or PlasmoAP, which had sensitivities of 89% and 84%, respectively, but outperformed the 40% sensitivity of ApicoAP (Fig 2C). All three algorithms as well as apicoplast BioID achieved high negative predictive values (NPV), since NPV is influenced by the larger number of true negatives (known non-apicoplast proteins) than true positives (known apicoplast) from literature data (S2A Fig). We expected that the advantages of apicoplast BioID would be its improved discrimination between true and false positives (Fig 2A) and the ability to detect proteins without classical targeting presequences. Indeed, bioinformatic algorithms had poor PPVs ranging from 19-36% compared to the 91% PPV of BioID (Fig 2D). Even a dataset consisting only of proteins predicted by all three algorithms achieved a PPV of just 25%. Similarly, the specificity of BioID outperformed that of the bioinformatic algorithms (S2B Fig). Consistent with the low PPVs of the bioinformatic algorithms, many proteins predicted by these algorithms are not enriched in BioID-Ap samples and are likely to be false positives (S3 Fig). Altogether, identification of apicoplast proteins using BioID provided a dramatic improvement in prediction performance over bioinformatic algorithms.

### Apicoplast BioID identifies proteins of diverse functions in multiple subcompartments

To determine whether lumenally targeted GFP-BirA* exhibited any labeling preferences, we assessed proteins identified based on the presence of transmembrane domains, their sub-organellar localization, and their functions. First, we determined the proportion of the 187 proteins identified by apicoplast BioID that are membrane proteins. To ensure that proteins were not classified as membrane proteins solely due to misclassification of a signal peptide as a transmembrane domain, we considered a protein to be in a membrane only if it contained at least one predicted transmembrane domain more than 80 amino acids from the protein’s *N*-terminus (as determined by annotation in PlasmoDB). These criteria suggested that 11% of identified proteins (20/187) were likely membrane proteins (Fig 3A), indicating that lumenal GFP-BirA* can label apicoplast membrane proteins.

**Fig 3.**
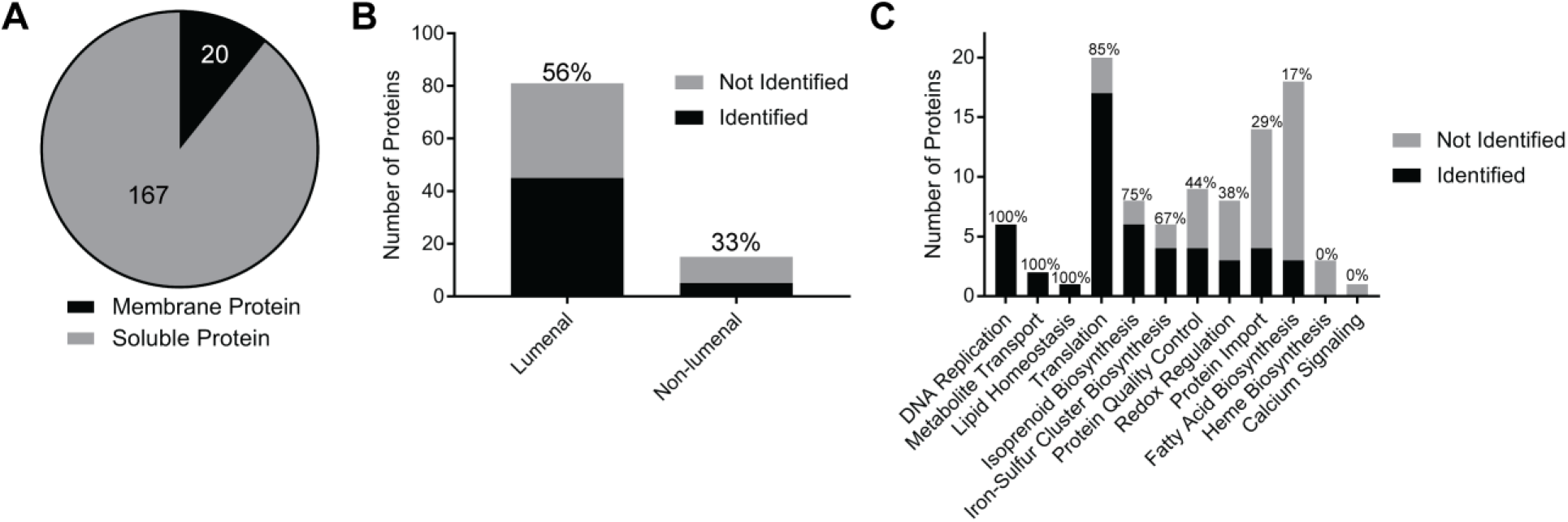
Diversity of protein labeling by apicoplast BioID. (A) Fraction of proteins identified by apicoplast BioID that are predicted to localize to a membrane. Proteins were considered “membrane” if they had at least one transmembrane domain annotated in PlasmoDB ending >80 amino acids from the annotated *N*-terminus. (B) Number of lumenal and non-lumenal positive controls identified. Percentages above bars indicate the percentage of known proteins from each category identified. (C) Number of proteins from established apicoplast pathways identified. Percentages above bars indicate the percentage of known proteins from each pathway identified.

Second, apicoplast proteins may localize to one or multiple sub-compartments defined by the four apicoplast membranes. It was unclear whether BirA* targeted to the lumen would label proteins in non-lumenal compartments. Based on literature descriptions, we classified the 96 known apicoplast proteins on our positive control list as either lumenal (present in lumenal space or on the innermost apicoplast membrane) or non-lumenal (all other sub-compartments) and determined the proportion that were identified in our dataset. Apicoplast BioID identified 56% (45/81) of the classified lumenal proteins and 33% (5/15) of the non-lumenal proteins (Fig 3B), suggesting that the GFP-BirA* bait used can label both lumenal and non-lumenal proteins but may have a preference for lumenal proteins (though this difference did not reach statistical significance).

Finally, we characterized the functions of proteins identified by apicoplast BioID. We grouped positive control apicoplast proteins into functional categories and assessed the proportion of proteins identified from each functional group (Fig 3C). BioID identified a substantial proportion (67-100%) of proteins in four apicoplast pathways that are essential in blood stage and localize to the apicoplast lumen, specifically DNA replication, protein translation, isoprenoid biosynthesis, and iron-sulfur cluster biosynthesis. Conversely, BioID identified few proteins involved in heme or fatty acid biosynthesis (0% and 17%, respectively), which are lumenal pathways that are non-essential in the blood-stage and which are likely to be more abundant in other life cycle stages [32–36]. We achieved moderate coverage of proteins involved in protein quality control (44%) and redox regulation (38%). Consistent with the reduced labeling of non-lumenal apicoplast proteins, only a small subset (29%) of proteins involved in import of nuclear-encoded apicoplast proteins were identified. Overall, apicoplast BioID identified soluble and membrane proteins of diverse functions in multiple apicoplast compartments with higher coverage for lumenal proteins required during blood-stage infection.

### The PlastNN algorithm expands the predicted apicoplast proteome with high accuracy

Apicoplast BioID provided the first experimental profile of the blood-stage apicoplast proteome but is potentially limited in sensitivity due to 1) difficulty in detecting low abundance peptides in complex mixtures; 2) inability of the promiscuous biotin ligase to access target proteins that are buried in membranes or protein complexes; or 3) stage-specific protein expression. Currently available bioinformatic predictions of apicoplast proteins circumvent these limitations, albeit at the expense of a low PPV (Fig 2D). We reasoned that increasing the number of high-confidence apicoplast proteins used to train algorithms could improve the accuracy of a prediction algorithm while maintaining high sensitivity. In addition, inclusion of exported proteins that traffic through the ER, which are common false positives in previous prediction algorithms, would also improve our negative training set.

We used our list of previously known apicoplast proteins (S2 Table) as well as newly-identified apicoplast proteins from BioID (S1 Table) to construct a positive training set of 205 apicoplast proteins (S4 Table). As a negative training set, we used our previous list of 451 signal peptide-containing non-apicoplast proteins (S2 Table). For each of the 656 proteins in the training set, we calculated the frequencies of all 20 canonical amino acids in a 50 amino acid region immediately following the predicted signal peptide cleavage site. In addition, given that apicoplast proteins have a characteristic transcriptional profile in blood-stage parasites [37] and that analysis of transcriptional profile has previously enabled identification of apicoplast proteins in the related apicomplexan *Toxoplasma gondii* [38], we obtained transcript levels at 8 time points during intraerythrocytic development from previous RNA-Seq data [39]. Altogether, each protein was represented by a vector of dimension 28 (20 amino acid frequencies plus 8 transcript levels). These 28-dimensional vectors were used as inputs to train a neural network with 3 hidden layers (Fig 4A and S5 Table). Six-fold cross-validation was used for training, wherein the training set was divided into 6 equal parts (folds) to train 6 separate models. Each time, 5 folds were used to train the model and 1 fold to measure the performance of the trained model.

**Fig 4.**
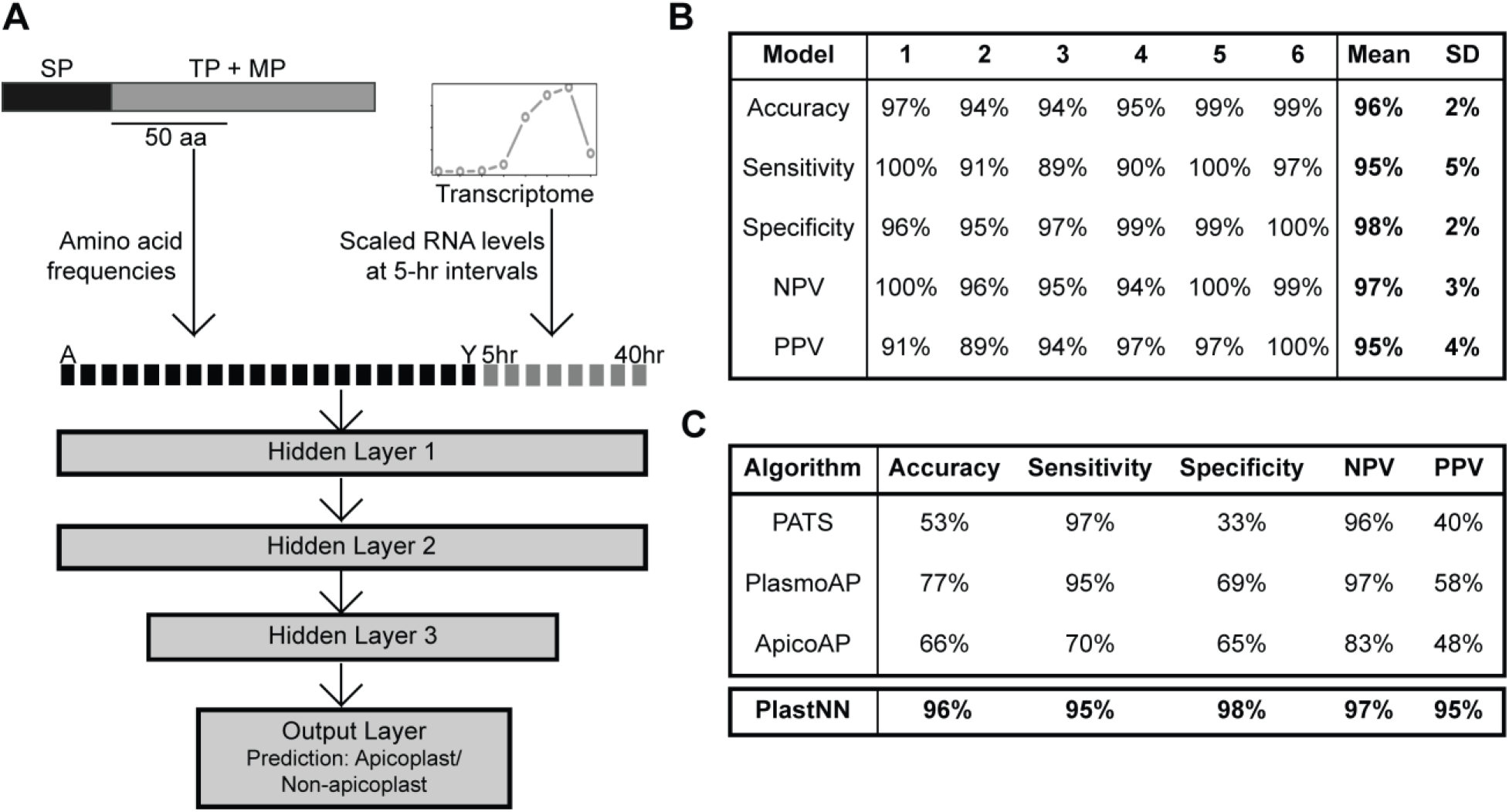
Improved prediction of apicoplast proteins using the PlastNN algorithm. (A) Schematic of the PlastNN algorithm. For each signal peptide-containing protein, a region of 50 amino acids immediately following the signal peptide cleavage site was selected and the frequencies of the 20 canonical amino acids in this region were calculated, resulting in a vector of length 20. Scaled RNA levels of the gene encoding the protein at 8 time points were added, resulting in a 28-dimensional vector representing each protein. This was used as input to train a neural network with 3 hidden layers, resulting in a prediction of whether the protein is targeted to the apicoplast or not. (B) Table showing the performance of the 6 models in PlastNN. Each model was trained on 5/6th of the training set and cross-validated on the remaining 1/6th. Values shown are accuracy, sensitivity, specificity, NPV, and PPV on the cross-validation set. The final values reported are the average and standard deviation over all 6 models. (C) Comparison of accuracy, sensitivity, specificity, NPV, and PPV for three previous algorithms and PlastNN.

We named this model PlastNN (ApicoPLAST Neural Network). PlastNN recognized apicoplast proteins with a cross-validation accuracy of 96 ± 2% (mean ± SD across 6 models), along with sensitivity of 95 ± 5% and PPV of 95 ± 4% (Fig 4B). This performance was higher than logistic regression on the same dataset (average accuracy = 91%; S6 Table). Combining the transcriptome features and the amino acid frequencies improves performance: the same neural network architecture with amino acid frequencies alone as input resulted in a lower average accuracy of 91%, while using transcriptome data alone resulted in an average accuracy of 90% (S6 Table). Comparison of the performance of PlastNN to existing prediction algorithms indicates that PlastNN distinguishes apicoplast and non-apicoplast proteins with higher accuracy than any previous prediction method (Fig 4C). To identify new apicoplast proteins, PlastNN was used to predict the apicoplast status of 450 predicted signal peptide-containing proteins that were not in our positive or negative training sets. Since PlastNN is composed of 6 models, we designated proteins as “apicoplast” if plastid localization was predicted by ≥4 of the 6 models.

PlastNN predicts 118 out of the 450 proteins to be targeted to the apicoplast (S7 Table). Combining these results with those from apicoplast BioID (S1 Table) and with experimental localization of proteins from the literature (S2 Table) yielded a compiled proteome of 346 putative nuclear-encoded apicoplast proteins (S8 Table).

### The apicoplast proteome contains novel and essential proteins

To determine whether candidate apicoplast proteins from this study have the potential to reveal unexplored parasite biology or are candidate antimalarial drug targets, we assessed the novelty and essentiality of the identified proteins. We found that substantial fractions of the BioID and PlastNN proteomes (49% and 71%, respectively) and 50% of the compiled apicoplast proteome represented proteins that could not be assigned to an established apicoplast pathway and therefore might be involved in novel organellar processes (Fig 5A). Furthermore, we identified orthologs of identified genes in the 150 genomes present in the OrthoMCL database [40]: 39% of the compiled apicoplast proteome were unique to apicomplexan parasites, with 58% of these proteins found only in *Plasmodium* spp. (Fig 5B). Of the 61% of proteins that were conserved outside of the Apicomplexa, we note that many of these contain conserved domains or are components of well-established pathways, such as DNA replication, translation, and metabolic pathways (S8 Table). This analysis indicates that many of the newly identified proteins are significantly divergent from proteins in their metazoan hosts.

**Fig 5.**
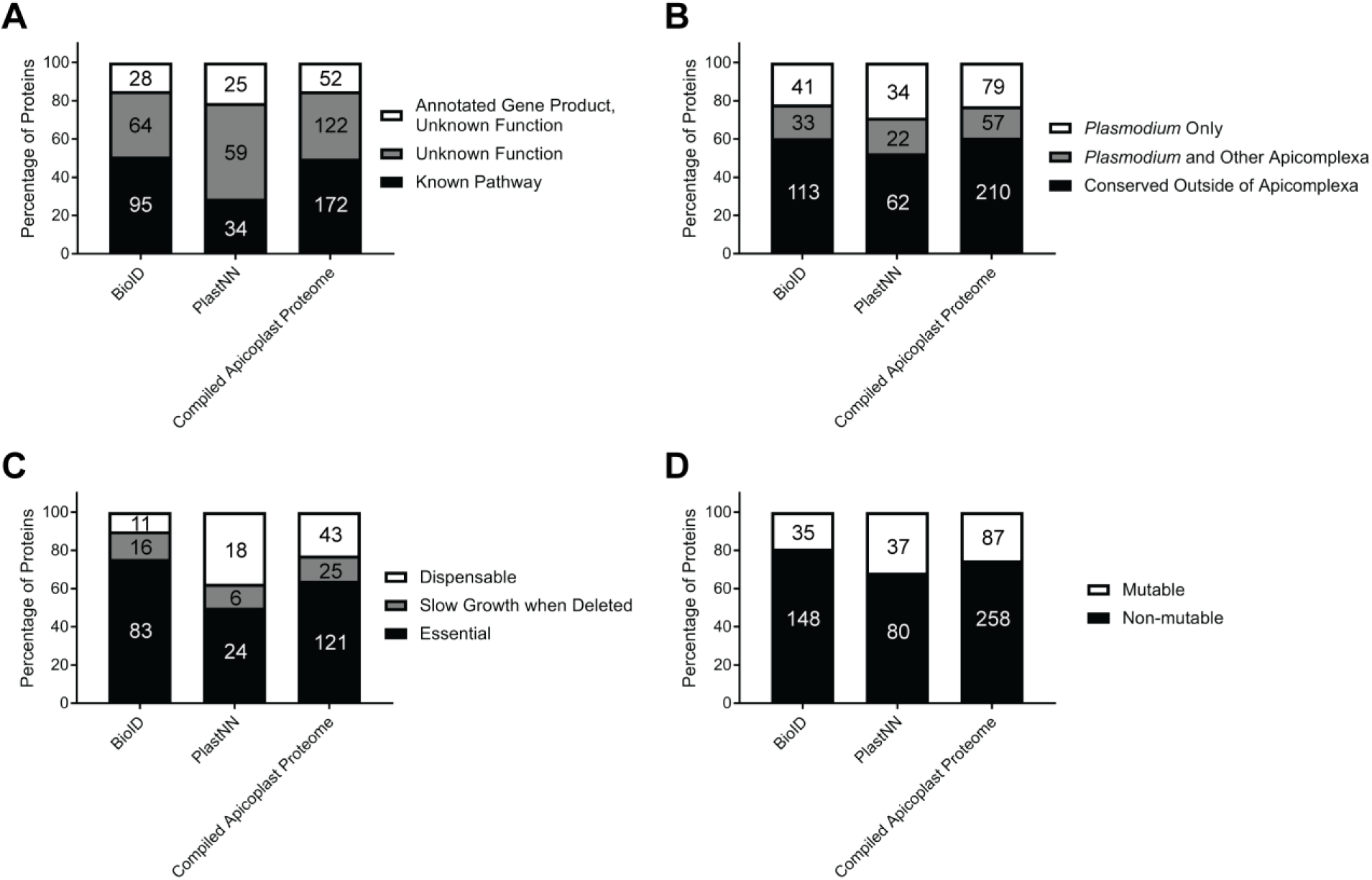
Apicoplast BioID identifies novel and essential proteins. (A) Percentage of proteins identified that have 1) annotated gene products but unknown function, 2) gene products annotated explicitly with “unknown function,” or 3) annotated gene products and function in a known cellular pathway. (B) Percentage of proteins identified that are *Plasmodium-* or Apicomplexa-specific based on OrthoMCL orthology analysis. (C) Percentage of proteins identified that are essential, cause slow growth when deleted, or are dispensable based on PlasmoGEM essentiality data of *P. berghei* orthologs [41]. (D) Percentage of proteins identified that were classified as mutable or non-mutable based on genome-scale transposon mutagenesis in *P. falciparum* [42]. In each panel, absolute numbers of proteins are indicated within bars.

Consistent with the critical role of the apicoplast in parasite biology, a genome-scale functional analysis of genes in the rodent malaria parasite *P. berghei* showed that numerous apicoplast proteins are essential for blood-stage survival [41]. Using this dataset, we found that 77% of proteins in the compiled apicoplast proteome that had *P. berghei* homologs analyzed by PlasmoGEM were important for normal blood-stage parasite growth (Fig 5C). Notably, of 49 proteins that were annotated explicitly with “unknown function” in their gene description and for which essentiality data are available, 38 are important for normal parasite growth, indicating that the high rate of essentiality for apicoplast proteins is true of both previously known and newly discovered proteins. In concordance with the PlasmoGEM data, recent genome-scale transposon mutagenesis in *P. falciparum* [42] identified 75% of proteins in the compiled apicoplast proteome as non-mutable (Fig 5D), suggesting essential functions in the blood stage. Overall, these data suggest that we have identified dozens of novel proteins that are likely critical for apicoplast biology.

### Localization of candidate apicoplast proteins identifies novel proteins of biological interest

Our analyses of the apicoplast BioID and PlastNN datasets suggested that these approaches enabled accurate, large-scale identification of apicoplast proteins (Figs 2D and 4C) and included many proteins of potential biological interest due to their novelty or their essentiality in the blood stage (Fig 5). As proof-of-concept of the utility of these datasets, several newly identified apicoplast proteins were experimentally validated. Fortuitously, while this manuscript was in preparation, 7 new apicoplast membrane proteins in *P. berghei* were validated by Sayers et al. [43]. Of these, apicoplast BioID identified the *P. falciparum* homologs of 3 proteins (PF3D7_1145500/ABCB3, PF3D7_0302600/ABCB4, and PF3D7_1021300) and PlastNN identified one (PF3D7_0908100). In addition to these, we also selected 4 candidates from apicoplast BioID and 2 from PlastNN to validate.

From the BioID list (S1 Table), we chose a rhomboid protease homolog ROM7 (PF3D7_1358300) and 3 conserved *Plasmodium* proteins of unknown function (PF3D7_0521400, PF3D7_1472800, and PF3D7_0721100) and generated cell lines expressing C-terminal GFP fusions from an ectopic locus in Dd2^attB^ parasites. With the exception of ROM7, which was chosen because of the biological interest of rhomboid proteases, we focused on proteins of unknown function to begin characterizing the large number of unannotated proteins in the *Plasmodium* genome (see Materials and Methods for additional candidate selection criteria).

To assess the apicoplast localization of each candidate, we first detected the apicoplast-dependent cleavage of each protein as a marker of its import. Most nuclear-encoded apicoplast proteins are proteolytically processed to remove *N*-terminal targeting sequences following successful import into the apicoplast [44, 45]. This processing is abolished in parasites rendered “apicoplast-minus” by treatment with an inhibitor (actinonin) to cause apicoplast loss [16, 46]. Comparison of protein molecular weight in apicoplast-intact and -minus parasites showed that ROM7, PF3D7_1472800, and PF3D7_0521400 (but not PF3D7_0721100) were cleaved in an apicoplast-dependent manner (Fig 6A).

**Fig 6.**
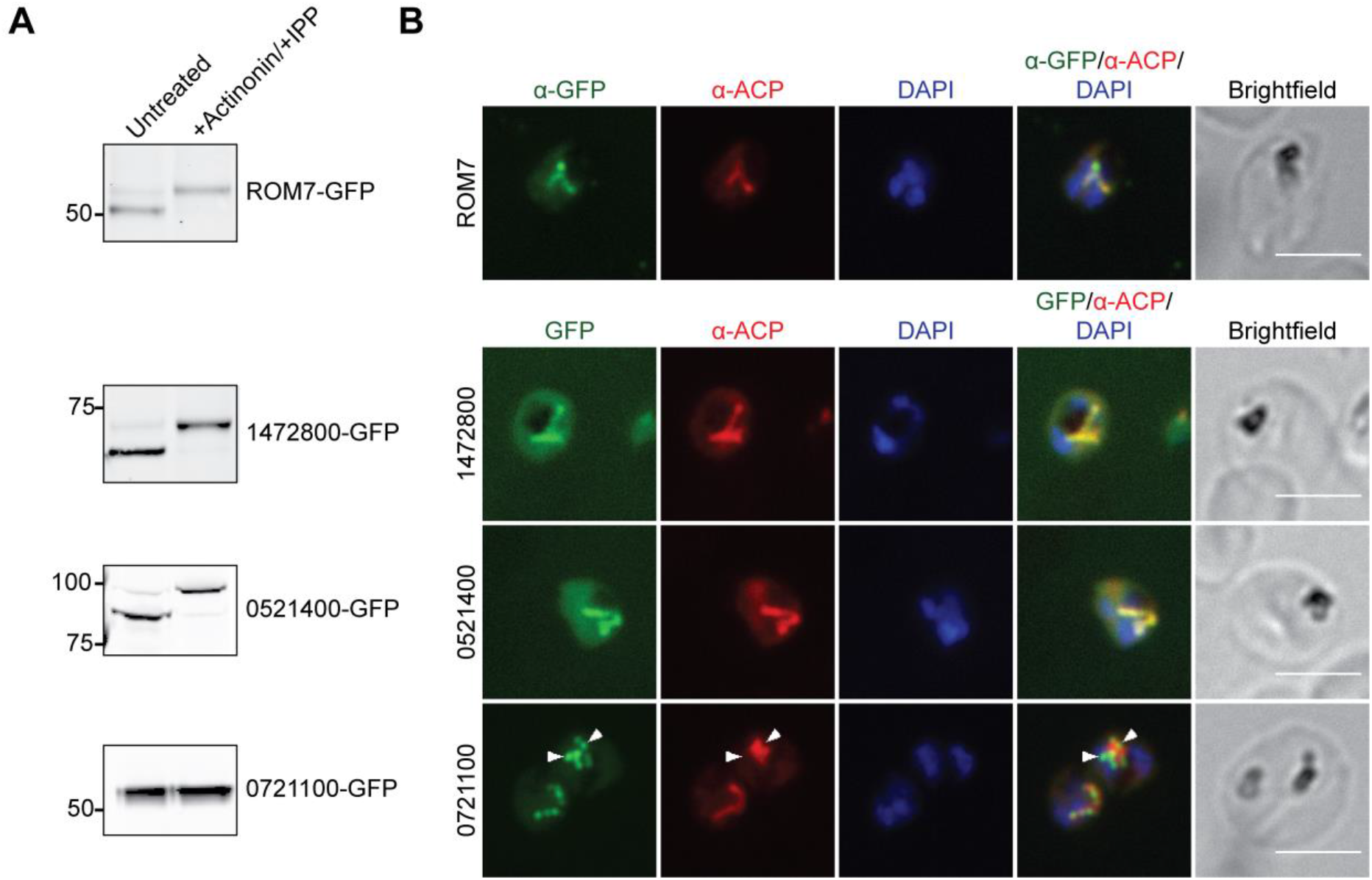
Localization of candidate apicoplast proteins identified by BioID. (A) Transit peptide processing assay for *C*-terminally GFP-tagged candidates. Ring-stage parasites were either untreated or treated with 10 μM actinonin/200 μM IPP for 3 days and protein processing was assessed by western blot. (B) Fixed-cell imaging of GFP-tagged candidates in parasites stained with an antibody raised against the apicoplast marker ACP. ROM7-GFP-expressing parasites were also stained with anti-GFP antibody due to low signal from intrinsic GFP fluorescence in fixed cells. Arrowheads indicate regions where PF3D7_0721100-GFP puncta appear adjacent to as opposed to co-localizing with ACP. Scale bars, 5 μm.

Next, we demonstrated co-localization of these three proteins with the apicoplast marker ACP by co-immunofluorescence analysis (co-IFAs; Fig 6B). ROM7, PF3D7_1472800, and PF3D7_0521400 clearly co-localized with ACP. PF3D7_0721100 localized to few large puncta not previously described for any apicoplast protein, which partly co-localized with the apicoplast marker ACP (Fig 6B and S4 Fig) but also appeared adjacent to ACP staining (Fig 6B and S4 Fig, arrowheads).

Finally, we localized the candidate-GFP fusions by live fluorescence microscopy and assessed the apicoplast dependence of their localization. ROM7-GFP, PF3D7_1472800-GFP, and PF3D7_0521400-GFP localized to branched structures characteristic of the apicoplast (S5 Fig). Upon actinonin treatment to render parasites “apicoplast-minus,” these proteins mislocalized to diffuse puncta (S5 Fig) previously observed for known apicoplast proteins [46]. Interestingly, while in untreated live parasites PF3D7_0721100-GFP again localized to a few large bright puncta, this protein also relocalized to the typical numerous diffuse puncta seen for genuine apicoplast proteins in apicoplast-minus parasites (S5 Fig).

Taken together, these data validate the apicoplast localization of ROM7, PF3D7_0521400, and PF3D7_1472800. Though transit peptide cleavage and the characteristic branched structure were not detected for PF3D7_0721100, partial co-localization with ACP and the mislocalization of PF3D7_0721100-GFP to puncta characteristic of apicoplast-minus parasites indicates that this protein may also be a true apicoplast protein. Further studies using either endogenously tagged protein or antibody raised against endogenous protein will be necessary to better characterize this localization.

From the PlastNN list (S7 Table), we selected 2 proteins of unknown function, PF3D7_1349900 and PF3D7_1330100. As above, each protein was appended with a C-terminal GFP tag and expressed as a second copy in Dd2^attB^ parasites. In agreement with apicoplast localization for each of these proteins, actinonin-mediated apicoplast loss caused loss of transit peptide processing (Fig 7A) and redistribution from a branched structure to diffuse puncta (S6 Fig). Furthermore, both proteins co-localized with the apicoplast marker ACP (Fig 7B).

**Fig 7.**
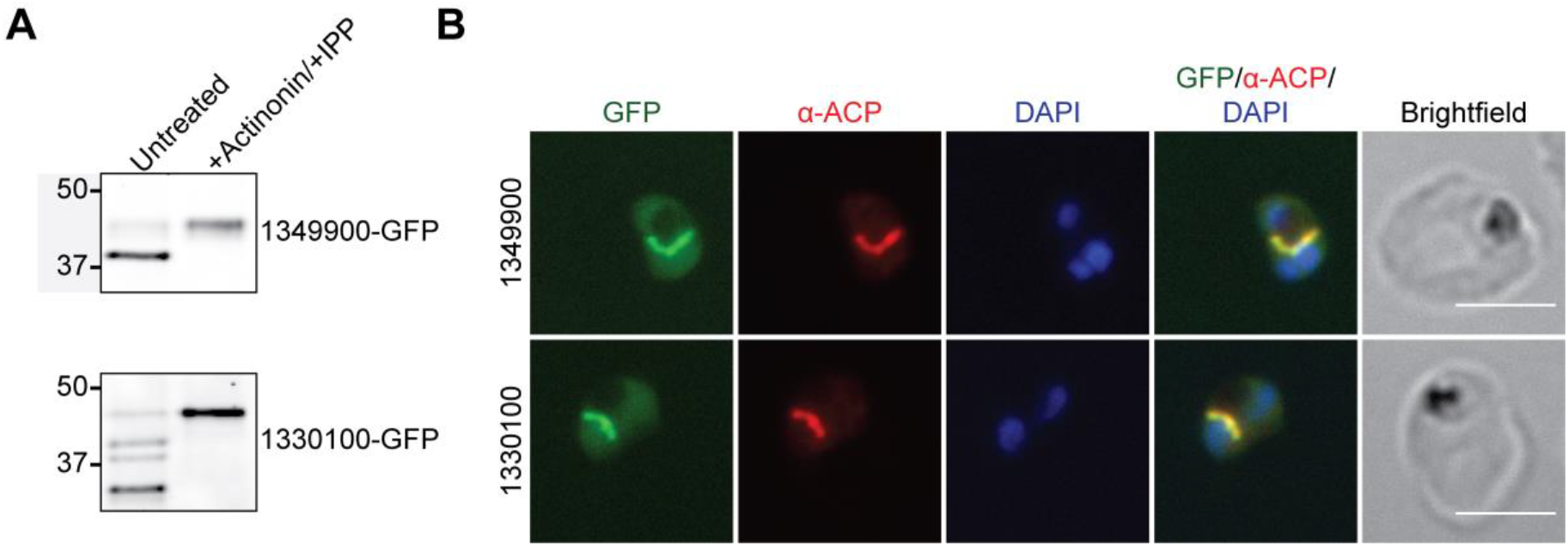
Localization of candidate apicoplast proteins identified by PlastNN. (A) Transit peptide processing assay for *C*-terminally GFP-tagged candidates. Ring-stage parasites were either untreated or treated with 10 μM actinonin/200 μM IPP for 3 days and protein processing was assessed by western blot. (B) Fixed-cell imaging of GFP-tagged candidates in parasites stained with an antibody raised against the apicoplast marker ACP. Scale bars, 5 μm.

Altogether, we confirmed the apicoplast localization of 5 novel apicoplast proteins, with a sixth protein (PF3D7_0721100) having potential apicoplast localization. These results, combined with validation of 4 apicoplast membrane proteins predicted in our datasets by Sayers et al., show that the apicoplast BioID and PlastNN datasets can successfully be used to prioritize apicoplast proteins of biological interest.

### A novel apicoplast protein ABCF1 is essential and required for organelle biogenesis

Given the potential of ATP binding cassette (ABC) proteins as drug targets, we sought to experimentally validate the essentiality of newly discovered apicoplast ABC proteins and assess their roles in metabolism or organelle biogenesis. Apicoplast BioID identified four ABC proteins: 3 ABCB-family proteins (ABCB3, ABCB4, and ABCB7) and an ABCF-family protein (ABCF1). We expected that these proteins might be important for apicoplast biology, as ABCB-family proteins are integral membrane proteins that typically act as small molecule transporters and ABCF-family proteins, which do not contain transmembrane domains, are typically involved in translation regulation [47, 48]. We pursued reverse genetic characterization of ABCB7 (PF3D7_1209900) and ABCF1 (PF3D7_0813700), as the essentiality of ABCB3 and ABCB4 has been previously studied [43, 49].

To assess localization and function of ABCB7 and ABCF1, we modified their endogenous loci to contain a *C*-terminal triple HA tag and tandem copies of a tetracycline repressor (TetR)-binding RNA aptamer in the 3’ UTR of either gene (S7 Fig) [50, 51]. Co-IFA confirmed ABCF1-3xHA colocalization with the apicoplast marker ACP (Fig 8A). ABCB7-3xHA localized to elongated structures that may be indicative of an intracellular organelle but rarely co-localized with ACP, indicating that it has a primarily non-apicoplast localization and is likely a false positive from the BioID dataset (S8A Fig).

**Fig 8.**
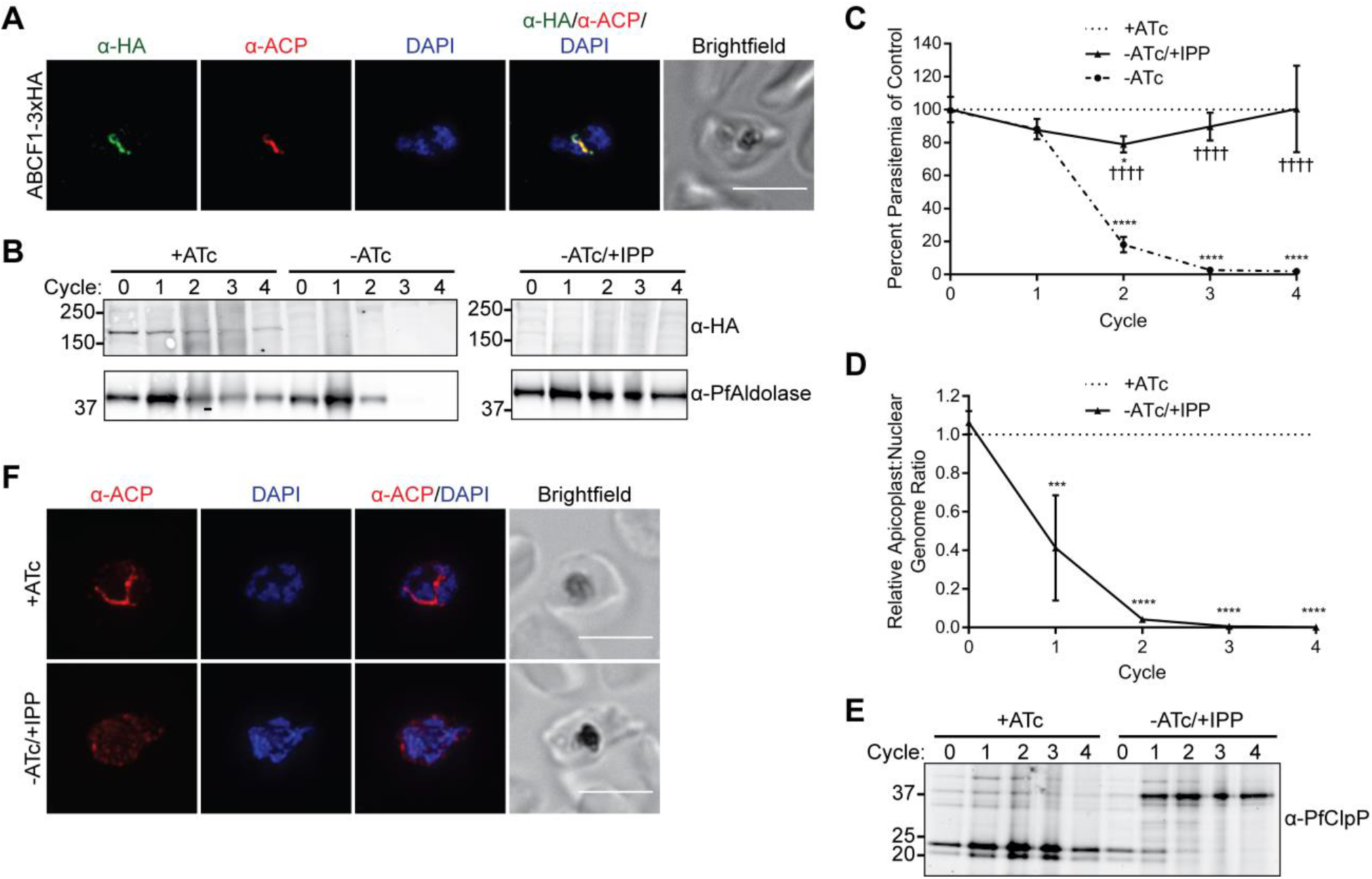
ABCF1 is an essential apicoplast protein required for organelle biogenesis. (A) Fixedcell imaging of ABCF1-3xHA knockdown parasites stained with antibodies raised against the HA tag and the apicoplast marker ACP. Scale bar, 5 μm. (B-F) ABCF1-3xHA knockdown parasites were grown in the presence of ATc (+ATc), the absence of ATc (-ATc), or the absence of ATc with IPP supplementation (-ATc/+IPP) for 4 growth cycles. (B) Western blot of ABCF1-3xHA expression. (C) Parasite growth. At each time point, data are normalized to the untreated (+ATc) control. Error bars represent standard deviation from the mean of two biological replicates. \**P* < 0.05, \*\*\*\**P* < 0.0001 compared to untreated control, ††††*P* < 0.0001 compared to -ATc condition, repeated measures two-way ANOVA with Tukey’s multiple comparisons test. (D) Apicoplast:nuclear genome ratio. At each time point, data are normalized to the untreated (+ATc) control. Error bars represent standard deviation from the mean of two biological replicates, each performed in technical triplicate. \*\*\**P* < 0.001, \*\*\*\**P* < 0.0001, repeated measures two-way ANOVA with Sidak’s multiple comparisons test. (E) Western blot of ClpP processing. (F) Fixed-cell imaging showing ACP localization after 2 cycles of knockdown. Scale bars, 5 μm.

Taking advantage of the TetR-binding aptamers in the 3’ UTR of ABCF1, we determined the essentiality and knockdown phenotype of this protein. In the presence of anhydrotetracycline (ATc), binding of the aptamer by a TetR-DOZI repressor is inhibited and ABCF1 is expressed. Upon removal of ATc, repressor binding blocks gene expression [50, 51]. Knockdown of ABCF1 caused robust parasite growth inhibition (Fig 8B-C). Growth inhibition of ABCF1-deficient parasites was reversed in the presence of isopentenyl pyrophosphate (IPP) (Fig 8C), which bypasses the need for a functional apicoplast [46], indicating that ABCF1 has an essential apicoplast function. Essential apicoplast functions can be placed into two broad categories: those involved in organelle biogenesis, and those involved solely in IPP production. Disruption of proteins required for organelle biogenesis causes apicoplast loss, while disruption of proteins involved in IPP production does not [16, 46, 52]. We determined whether knockdown of ABCF1 caused apicoplast loss by assessing 1) absence of the apicoplast genome, 2) loss of transit peptide processing of nuclear-encoded apicoplast proteins, and 3) relocalization of apicoplast proteins to puncta. Indeed, the apicoplast:nuclear genome ratio drastically decreased in ABCF1 knockdown parasites beginning 1 cycle after knockdown (Fig 8D), and western blot showed that the apicoplast protein ClpP was not processed in ABCF1 knockdown parasites (Fig 8E).

Furthermore, IFA of the apicoplast marker ACP confirmed redistribution from an intact plastid to diffuse cytosolic puncta (Fig 8F). In contrast to ABCF1, a similar knockdown of ABCB7 caused no observable growth defect after four growth cycles despite significant reduction in protein levels (S8B-C Fig). Together, these results show that ABCF1 is a novel and essential apicoplast protein with a previously unknown function in organelle biogenesis.

## Discussion

Since the discovery of the apicoplast, identification of its proteome has been a pressing priority. We report the first large-scale proteomic analysis of the apicoplast in blood-stage malaria parasites, which identified 187 candidate proteins with 52% sensitivity and 91% PPV. A number of groups have also profiled parasite-specific membrane compartments using proximity biotinylation but observed contamination with proteins in or trafficking through the ER, preventing accurate identification of these proteomes without substantial manual curation and validation [23, 24, 26–29]. This background labeling is expected since proteins traffic through the ER en route to several parasite-specific compartments, including the parasitophorous vacuole, host cytoplasm, food vacuole, and invasion organelles. The high specificity of our apicoplast BioID proteome depended on 1) the use of a control cell line expressing ER-localized GFP-BirA* to detect enrichment of apicoplast proteins from background ER labeling and 2) strong positive and negative controls to set an accurate threshold. We suspect a similar strategy to detect nonspecific ER background may also improve the specificity of proteomic datasets for other parasite-specific, endomembrane-derived compartments.

Leveraging our successful proteomic analysis, we used these empirical data as an updated training set to also improve computational predictions of apicoplast proteins. PlastNN identified an additional 118 proteins with 95% sensitivity and 95% PPV. Although two previous prediction algorithms, PATS and ApicoAP, also applied machine learning to the problem of transit peptide prediction, we reasoned that their low accuracy arose from the small training sets used (ApicoAP) and the use of cytosolic as well as endomembrane proteins in the negative training set (PATS). By using an expanded positive training set based on proteomic data and limiting our training sets to only signal peptide-containing proteins, we developed an algorithm with higher sensitivity than BioID and higher accuracy than previous apicoplast protein prediction models. Inevitably some false positives from the BioID dataset would have been used for neural network training and cross-validation. While this may slightly influence the PPV of the PlastNN list, we expect that the substantially larger fraction of true positives in the training set mitigated the effects of any false positives. Importantly, as more apicoplast and non-apicoplast proteins in *P. falciparum* parasites are experimentally validated, updated training sets can be used to re-train PlastNN. Moreover, PlastNN suggests testable hypotheses regarding the contribution of sequence-based and temporal regulation to protein trafficking in the ER.

Overall, we have compiled a high-confidence apicoplast proteome of 346 proteins that are rich in novel and essential functions (Fig 5). This proteome likely represents a majority of soluble apicoplast proteins, since 1) our bait for proximity biotinylation targeted to the lumen and 2) most soluble proteins use canonical targeting sequences that can be predicted. An important next step will be to expand the coverage of apicoplast membrane proteins, which more often traffic via distinctive routes [53, 54]. Performing proximity biotinylation with additional bait proteins may identify such atypical apicoplast proteins. In the current study, our bait was an inert fluorescent protein targeted to the apicoplast lumen to minimize potential toxicity of the construct. The success of this apicoplast GFP bait gives us confidence to attempt more challenging baits, including proteins localized to sub-organellar membrane compartments or components of the protein import machinery. Performing apicoplast BioID in liver and mosquito stages may also define apicoplast functions in these stages. This compiled proteome represents a substantial improvement upon previous bioinformatics predictions of apicoplast proteins and provides a strong foundation for further refinement. In analogy to progress on the mammalian mitochondrial proteome, which over the course of decades has been expanded and refined by a combination of proteomic, computational, and candidate-based approaches [55, 56], we expect that future proteomic, computational, and candidate-based approaches to identify apicoplast proteins will be critical for ultimately determining a comprehensive apicoplast proteome,

Organellar proteomes are valuable hypothesis-generating tools. Already several candidates of biological interest based on their biochemical function annotations were validated. We demonstrated an unexpected role for the ATP-binding cassette protein PfABCF1 in apicoplast biogenesis. ABCF proteins are understudied compared to other ABC-containing proteins but tend to have roles in translation regulation [47]. An *E. coli* homolog, EttA, regulates translation initiation in response to cellular ATP levels [57, 58], and mammalian and yeast ABCF1 homologs also interact with ribosomes and regulate translation [59–62]. By analogy, PfABCF1 may regulate the prokaryotic translation machinery in the apicoplast, although the mechanistic basis for the severe defect in parasite replication upon loss of PfABCF1 is unclear.

We also validated *Pf*ROM7 as an apicoplast-localized rhomboid protease. Rhomboid proteases are a diverse family of intramembrane serine proteases found in all domains of life. In the Apicomplexa, rhomboids have been studied primarily for their roles in processing adhesins on the parasite cell surface [63], although the functions of most apicomplexan rhomboids are still unknown. Little is known about ROM7 other than that it appears to be absent from coccidians and was refractory to deletion in *P. berghei* [64, 65]. However, a rhomboid protease was recently identified as a component of symbiont-derived ERAD-like machinery (SELMA) that transports proteins across a novel secondary plastid membrane in diatoms [66], indicating that ROM7 may similarly play a role in apicoplast protein import in *Plasmodium* parasites. Neither PfABCF1 nor *Pf*ROM7 had known roles in the apicoplast prior to their identification in this study, underscoring the utility of unbiased approaches to identify new organellar proteins. Moreover, the apicoplast is one of few models for complex plastids that permits functional analysis of identified proteins to investigate the molecular mechanisms underpinning serial endosymbiosis. A summary of all candidate proteins validated in this study is shown in S9 Table.

A recent study aimed at identifying apicoplast membrane transporters highlights the difficulty in identifying novel apicoplast functions in the absence of a high-confidence proteome [43]. Taking advantage of the tractable genetics in murine *Plasmodium* species, Sayers et al. screened 27 candidates in *P. berghei* for essentiality and apicoplast localization. Following >50 transfections, 3 essential and 4 non-essential apicoplast membrane proteins were identified. One newly identified essential apicoplast membrane protein was then validated to be required for apicoplast biogenesis in *P. falciparum.* In contrast, even though our study was not optimized to identify membrane proteins, the combination of BioID and PlastNN identified 2 known apicoplast transporters, 4 of the new apicoplast membrane protein homologs, and 56 additional proteins predicted to contain at least one transmembrane domain. A focused screen of higher quality candidates in *P. falciparum* is likely to be more rapid and yield the most relevant biology. Our high-confidence apicoplast proteome will streamline these labor-intensive screens, focusing on strong candidates for downstream biological function elucidation. As methods for analyzing gene function in *P. falciparum* parasites continue to improve, this resource will become increasingly valuable for characterizing unknown organellar pathways.

## Materials and Methods

### Parasite growth

*Plasmodium falciparum* Dd2^attB^ [31] (MRA-843) were obtained from MR4. NF54^Cas9+T7 Polymerase^ parasites [67] were a gift from Jacquin Niles. Parasites were grown in human erythrocytes (2% hematocrit) obtained from the Stanford Blood Center in RPMI 1640 media (Gibco) supplemented with 0.25% Albumax II (Gibco), 2 g/L sodium bicarbonate, 0.1 mM hypoxanthine (Sigma), 25 mM HEPES, pH 7.4 (Sigma), and 50 μg/L gentamicin (Gold Biotechnology) at 37°C, 5% O2, and 5% CO2.

### Vector construction

Oligonucleotides were purchased from the Stanford Protein and Nucleic Acid facility or IDT. gBlocks were ordered from IDT. Molecular cloning was performed using In-Fusion cloning (Clontech) or Gibson Assembly (NEB). Primer and gBlock sequences are available in S10 Table.

To generate the plasmid pRL2-ACPL-GFP for targeting transgenes to the apicoplast, the first 55 amino acids from ACP were PCR amplified with primers MB015 and MB016 and were inserted in front of the GFP in the pRL2 backbone [68] via the AvrII/BsiWI sites. To generate pRL2-ACPL-GFP-BirA* for targeting a GFP-BirA* fusion to the apicoplast, GFP was amplified from pLN-ENR-GFP using primers MB087 and MB088 and BirA* was amplified from pcDNA3.1 mycBioID (Addgene 35700) [20] using primers MB089 and MB090. These inserts were simultaneously cloned into BsiWI/AflII-digested pRL2-ACPl-GFP to generate pRL2-ACPL-GFP-BirA*. To generate pRL2-SP-GFP-BirA*-SDEL for targeting GFP-BirA* to the ER, SP-GFP-BirA*-SDEL was PCR amplified from pRL2-ACPL-GFP-BirA* using primers MB093 and MB094 and was cloned into AvrII/AflII-digested pRL2-ACP_L_-GFP. For GFP-tagging to confirm localization of proteins identified by apicoplast BioID, full-length genes were amplified from parasite cDNA with primers as described in S10 Table and were cloned into the AvrII/BsiWI sites of pRL2-ACP_L_-GFP.

For CRISPR-Cas9-based editing of endogenous ABCB7 and ABCF1 loci, sgRNAs were designed using the eukaryotic CRISPR guide RNA/DNA design tool (http://grna.ctegd.uga.edu/). To generate a linear plasmid for CRISPR-Cas9-based editing, left homology regions were amplified with primers MB256 and MB257 (ABCB7) or MB260 and MB261 (ABCF1) and right homology regions were amplified with MB258 and MB259 (ABCB7) or MB262 and MB263 (ABCF1). For each gene, a gBlock containing the recoded coding sequence C-terminal of the CRISPR cut site and a triple HA tag was synthesized with appropriate overhangs for Gibson Assembly. This fragment and the appropriate left homology region were simultaneously cloned into the FseI/ApaI sites of the linear plasmid pSN054-V5. Next, the appropriate right homology region and a gBlock containing the sgRNA expression cassette were simultaneously cloned into the AscI/I-SceI sites of the resultant vectors to generate the plasmids pSN054-ABCB7-TetR-DOZI and pSN054-ABCF1-TetR-DOZI.

### Parasite transfection

Transfections were carried out using variations on the spontaneous uptake method [69, 70]. In the first variation, 100 μg of each plasmid was ethanol precipitated and resuspended in 30 μL sterile TE buffer and was added to 150 μL packed RBCs resuspended to a final volume of 400 μL in cytomix. The mixture was transferred to a 0.2 cm electroporation cuvette (Bio-Rad) and was electroporated at 310 V, 950 μF, infinity resistance in a Gene Pulser Xcell electroporation system (Bio-Rad) before allowing parasites to invade. Drug selection was initiated 3 days after transfection. Alternatively, 50 μg of each plasmid was ethanol precipitated and resuspended in 0.2 cm electroporation cuvettes in 100 μL TE buffer, 100 μL RPMI containing 10 mM HEPES-NaOH, pH 7.4, and 200 μL packed uninfected RBCs. RBCs were pulsed with 8 square wave pulses of 365 V x 1 ms separated by 0.1 s. RBCs were allowed to reseal for 1 hour in a 37°C water bath before allowing parasites to invade. Drug selection was initiated 4 days after transfection. All transfectants were selected with 2.5 μg/mL Blasticidin S (Research Products International). Additionally, BioID-ER parasites were selected with 125 μg/mL G418 sulfate (Corning) and ABCB7 and ABCF1 TetR-DOZI parasites were grown in the presence of 500 nM ATc. Transfections for generating BioID constructs (Fig 1) and expression of GFP-tagged candidates (Figs 6 and 7) were performed in the Dd2^attB^ background. Transfections for CRISPR editing were performed with the NF54^Cas9^+^T7 Polymerase^ background and clonal parasite lines were obtained by limiting dilution.

Correct modification of transfectant genomes was confirmed by PCR. Briefly, 200 μL of 2% hematocrit culture was pelleted and resuspended in water, and 2 μL of the resulting lysate was used as template for PCR with Phusion polymerase (NEB). PCR targets and their corresponding primer pairs are as follows: integrated *attL* site, p1 + p2; integrated *attR* site, MW001 + MW003; unintegrated *attB* site, MW004 + MW003; ABCB7 unintegrated left homology region (LHR), MB269 + MB270; ABCB7 integrated LHR, MB269 + MB255;

ABCB7 unintegrated right homology region (RHR), MB281 + MB278; ABCB7 integrated RHR, MB276 + MB278; ABCF1 unintegrated LHR, MB271 + MB272; ABCF1 integrated LHR, MB271 + MB255; ABCF1 unintegrated RHR, MB282 + MB283; ABCF1 integrated RHR, MB276 + MB283.

### Biotin labeling

To label parasites for analysis by streptavidin blot, fixed imaging, or mass spectrometry, cultures of majority ring-stage parasites were treated with 50 μM biotin or with a DMSO vehicle-only control. Cultures were harvested for analysis 16 hours later as majority trophozoites and schizonts.

### Actinonin treatment and IPP rescue

To generate apicoplast-minus parasites, ring-stage cultures were treated with 10 μM actinonin (Sigma) and 200 μM IPP (Isoprenoids, LLC) and cultured for 3 days before analysis.

### Western blotting

Parasites were separated from RBCs by lysis in 0.1% saponin and were washed in PBS. Parasite pellets were resuspended in PBS containing 1X NuPAGE LDS sample buffer with 50 mM DTT and were boiled at 95°C for 10 minutes before separation on NuPAGE or Bolt Bis-Tris gels and transfer to nitrocellulose. Membranes were blocked in 0.1% Hammarsten casein (Affymetrix) in 0.2X PBS with 0.01% sodium azide. Antibody incubations were performed in a 1:1 mixture of blocking buffer and TBST (Tris-buffered saline with Tween-20; 10 mM Tris, pH 8.0, 150 mM NaCl, 0.25 mM EDTA, 0.05% Tween 20). Blots were incubated with primary antibody for either 1 hour at room temperature or at 4°C overnight at the following dilutions: 1:20,000 mouse-α-GFP JL-8 (Clontech 632381); 1:20,000 *rabbit-α-Plasmodium* aldolase (Abcam ab207494); 1:1000 rat-α-HA 3F10 (Sigma 11867423001); 1:4000 rabbit-α-*Pf*ClpP [71]. Blots were washed once in TBST and were incubated for 1 hour at room temperature in a 1:10,000 dilution of the appropriate secondary antibody: IRDye 800CW donkey-α-rabbit; IRDye 680LT goat-α-mouse; IRDye 680LT goat-α-rat (LI-COR Biosciences). For detection of biotinylated proteins, blots were incubated with 1:1000 IRDye 680RD streptavidin for one hour at room temperature. Blots were washed three times in TBST and once in PBS before imaging on a LI-COR Odyssey imager.

### Microscopy

For live imaging, parasites were settled onto glass-bottom microwell dishes (MatTek P35G-1.5-14-C) or Lab-Tek II chambered coverglass (ThermoFisher 155409) in PBS containing 0.4% glucose and 2 μg/mL Hoechst 33342 stain (ThermoFisher H3570).

For fixed imaging of biotinylated proteins in cells, biotin-labeled parasites were processed as in Tonkin et al. [72] with modifications. Briefly, parasites were washed in PBS and were fixed in 4% paraformaldehyde (Electron Microscopy Science 15710) and 0.015% glutaraldehyde (Electron Microscopy Sciences 16019) in PBS for 30 minutes. Cells were washed once in PBS, resuspended in PBS, and allowed to settle onto poly-L-lysine-coated coverslips (Corning) for 60 minutes. Coverslips were then washed once with PBS, permeabilized in 0.1% Triton X-100 in PBS for 10 minutes, and washed twice more in PBS. Cells were treated with 0.1 mg/mL sodium borohydride in PBS for 10 minutes, washed once in PBS, and blocked in 3% BSA in PBS. To visualize biotin-labeled proteins, coverslips were incubated with 1:1000 AlexaFluor 546-conjugated streptavidin (ThermoFisher S11225) for one hour followed by three washes in PBS. No labeling of GFP was necessary, as these fixation conditions preserve intrinsic GFP fluorescence [72]. Coverslips were mounted onto slides with ProLong Gold antifade reagent with DAPI (ThermoFisher) and were sealed with nail polish prior to imaging.

For immunofluorescence analysis, parasites were processed as above except that fixation was performed with 4% paraformaldehyde and 0.0075% glutaraldehyde in PBS for 20 minutes and blocking was performed with 5% BSA in PBS. Following blocking, primary antibodies were used in 5% BSA in PBS at the following concentrations: 1:500 rabbit-α-PfACP [73]; 1:1000 rabbit-α-*Pf*Bip1:1000 (a gift from Sebastian Mikolajczak and Stefan Kappe); 1:500 mouse-α-GFP JL-8 (Clontech 632381); 1:100 rat-α-HA 3F10 (Sigma 11867423001). Coverslips were washed three times in PBS, incubated with goat-α-rat 488 (ThermoFisher A-11006), goat-α-mouse 488 (ThermoFisher A11029), or donkey-α-rabbit 568 (ThermoFisher A10042) secondary antibodies at 1:3000, and washed three times in PBS prior to mounting as above.

Live and fixed cells were imaged with 100X, 1.4 NA or 100X, 1.35 NA objectives on an Olympus IX70 microscope with a DeltaVision system (Applied Precision) controlled with SoftWorx version 4.1.0 and equipped with a CoolSnap-HQ CCD camera (Photometrics). Images were taken in a single z-plane, with the exception of those presented in Figs 1D, 8A, and S8A, which were captured as a series of z-stacks separated by 0.2 μm intervals, deconvolved, and displayed as maximum intensity projections. Brightness and contrast were adjusted in Fiji (ImageJ) for display purposes. Image capture and processing conditions were identical for micrographs of the same cell line when multiple examples are displayed (S4 Fig) or when comparing untreated to actinonin-treated cells (S5 and S6 Figs).

### Biotin pulldowns, mass spectrometry, and data analysis

Biotin-labeled parasites were harvested by centrifugation and were released from the host RBC by treatment with 0.1% saponin/PBS. Parasites were washed twice more with 0.1% saponin/PBS followed by PBS and were either used immediately for analysis or were stored at −80°C. Parasite pellets were resuspended in RIPA buffer [50 mM Tris-HCl, pH 7.4, 150 mM NaCl, 0.1% SDS, 0. 5% sodium deoxycholate, 1% Triton X-100, 1 mM EDTA] containing a protease inhibitor cocktail (Pierce) and were lysed on ice for 30 minutes with occasional pipetting. Insoluble debris was removed by centrifugation at 16,000 xg for 15 minutes at 4°C. Biotinylated proteins were captured using High Capacity Streptavidin Agarose beads (Pierce) for 2 hours at room temperature. Beads were then washed three times with RIPA buffer, three times with SDS wash buffer [50 mM Tris-HCl, pH 7.4, 150 mM NaCl, 2% SDS], six times with urea wash buffer [50 mM Tris-HCl, pH 7.4, 150 mM NaCl, 8 M urea], and three times with 100 mM ammonium bicarbonate. Proteins were reduced with 5 mM DTT for 60 minutes at 37°C followed by treatment with 14 mM iodoacetamide (Pierce) at room temperature for 45 minutes. Beads were washed once with 100 mM ammonium bicarbonate and were digested with 10 μg/mL trypsin (Promega) at 37°C overnight. The following day, samples were digested with an additional 5 μg/mL trypsin for 3-4 hours. Digested peptides were separated from beads by addition of either 35% or 50% final concentration acetonitrile, and peptides were dried on a SpeedVac prior to desalting with C18 stage tips.

Desalted peptides were resuspended in 0.1% formic acid and analyzed by online capillary nanoLC-MS/MS. Samples were separated on an in-house made 20 cm reversed phase column (100 μm inner diameter, packed with ReproSil-Pur C18-AQ 3.0 μm resin (Dr. Maisch GmbH)) equipped with a laser-pulled nanoelectrospray emitter tip. Peptides were eluted at a flow rate of 400 nL/min using a two-step linear gradient including 2-25% buffer B in 70 min and 25-40% B in 20 min (buffer A: 0.2% formic acid and 5% DMSO in water; buffer B: 0.2% formic acid and 5% DMSO in acetonitrile) in an Eksigent ekspert nanoLC-425 system (AB Sciex). Peptides were then analyzed using a LTQ Orbitrap Elite mass spectrometer (Thermo Scientific). Data acquisition was executed in data dependent mode with full MS scans acquired in the Orbitrap mass analyzer with a resolution of 60000 and m/z scan range of 340-1600. The top 20 most abundant ions with intensity threshold above 500 counts and charge states 2 and above were selected for fragmentation using collision-induced dissociation (CID) with isolation window of 2 m/z, normalized collision energy of 35%, activation Q of 0.25 and activation time of 5 ms. The CID fragments were analyzed in the ion trap with rapid scan rate. In additional runs, the top 10 most abundant ions with intensity threshold above 500 counts and charge states 2 and above were selected for fragmentation using higher energy collisional dissociation (HCD) with isolation window of 2 m/z, normalized collision energy of 35%, and activation time of 25 ms.

The HCD fragments were analyzed in the Orbitrap with a resolution of 15000. Dynamic exclusion was enabled with repeat count of 1 and exclusion duration of 30 s. The AGC target was set to 1000000, 50000, and 5000 for full FTMS scans, FTMSn scans and ITMSn scans, respectively. The maximum injection time was set to 250 ms, 250 ms, and 100 ms for full FTMS scans, FTMSn scans and ITMSn scans, respectively.

The resulting spectra were searched against a “target-decoy” sequence database [74] consisting of the PlasmoDB protein database (release 32, released April 19, 2017), the Uniprot human database (released February 2, 2015), and the corresponding reversed sequences using the SEQUEST algorithm (version 28, revision 12). The parent mass tolerance was set to 50 ppm and the fragment mass tolerance to 0.6 Da for CID scans, 0.02 Da for HCD scans. Enzyme specificity was set to trypsin. Oxidation of methionines was set as variable modification and carbamidomethylation of cysteines was set as static modification. Peptide identifications were filtered to a 1% peptide false discovery rate using a linear discriminator analysis [75]. Precursor peak areas were calculated for protein quantification.

### Apicoplast protein prediction algorithms and positive/negative control apicoplast proteins

To generate updated lists of PATS-predicted apicoplast proteins, nuclear-encoded *P. falciparum* 3D7 proteins (excluding pseudogenes) from PlasmoDB version 28 (released March 30, 2016) were used to check for existence of a putative bipartite apicoplast targeting presequence using the artificial neural network predictor PATS [17].

Updated PlasmoAP-predicted apicoplast proteins were identified using the PlasmoDB version 32 proteome (released April 19, 2017) by first checking for the presequence of a predicted signal peptide using the neural network version of SignalP version 3.0 [76], and were considered positive if they had a D-score above the default cutoff. The SignalP C-score was used to predict the signal peptide cleavage position, and the remaining portion of the protein was inspected for presence of a putative apicoplast transit peptide using the rules described for PlasmoAP [18], implemented in a Perl script.

*P. falciparum* proteins predicted to localize to the apicoplast by ApicoAP were accessed from the original paper [19]. Genes predicted to encode pseudogenes were excluded.

A positive control list of 96 high-confidence apicoplast proteins (S2 Table) was generated based on either (1) published localization of that protein in *Plasmodium* parasites or *Toxoplasma gondii* or (2) presence of that protein in either the isoprenoid biosynthesis or fatty acid biosynthesis/utilization pathways. To generate a negative control list of potential false positives, nuclear-encoded proteins (excluding pseudogenes) predicted to contain a signal peptide were identified as above and 451 of these proteins were designated as negative controls based on GO terms, annotations, and the published literature.

### Feature extraction for neural network

To generate the positive training set for PlastNN, we took the combined list of previously known apicoplast proteins (S2 Table) and apicoplast proteins identified by BioID (S1 Table) and removed proteins that (1) were likely false positives based on our negative control list (S2 Table) or published localization data; (2) were likely targeted to the apicoplast without the canonical bipartite *N*-terminal leader sequence; or (3) did not contain a predicted signal peptide based on the SignalP 3.0 D-score. This yielded a final positive training set of 205 proteins (S4 Table). The negative training set was the previously generated list of known non-apicoplast proteins (S2 Table). The test set for PlastNN consisted of 450 proteins predicted to have a signal peptide by the SignalP 3.0 D-score that were not in the positive or negative training sets.

For each protein in our training and test sets, we took the 50 amino acids immediately after the end of the predicted signal peptide (according to the SignalP 3.0 C-score) and calculated the frequency of each of the 20 amino acids in this sequence. The length of 50 amino acids was chosen empirically by trying lengths from 20-100; highest accuracy was obtained using 50. Scaled FPKM values at 8 time points during intraerythrocytic development were obtained from published RNA-Seq [39]. By combining the amino acid frequencies with the 8 transcriptome values, we represented each protein in our training and test sets by a feature vector of length 28.

### Neural network training and cross-validation

To train the model, the 205 positive and 451 negative training examples were combined and randomly shuffled. The training set was divided into 6 equal folds, each containing 109 or 110 examples. We trained models using 6-fold cross-validation; that is, we trained 6 separate models with the same architecture, each using 5 of the 6 folds for training and then using the one remaining fold as a cross-validation set to evaluate performance. Accuracy, sensitivity, specificity, NPV, and PPV are calculated on this cross-validation set. The final reported values of accuracy, sensitivity, specificity, NPV, and PPV are the average and standard deviation over all 6 models. When predicting on the test set, the final predictions are generated by a majority vote of all 6 models.

Neural networks were trained using the RMSProp optimization algorithm with a learning rate of 0.0001. Tensorflow version 1.4.1 was used to build and train the neural network. Logistic regression on the same dataset was carried out using the caret package (version 6.0-77) in R version 3.3.3.

### Analyses of Apicoplast Proteome Datasets

The BioID apicoplast proteome and the predicted proteomes from PATS, PlasmoAP, ApicoAP, and PlastNN were analyzed according to the following formulae:

Accuracy = (TP + TN)/(TP + FP + TN + FN)

Sensitivity = TP/(TP + FN)

Specificity = TN/(TN + FP)

Negative Predictive Value (NPV) = TN/(TN + FN)

Positive Predictive Value (PPV) = TP/(TP + FP)

Abbreviations: TP, true positive; TN, true negative; FP, false positive; FN, false negative. Because none of the 451 negative control proteins from the original list (S2 Table) were identified in our 187-protein BioID proteome, we manually inspected the BioID list, identified 5 likely false positives, and added these to the negative control list for the purposes of analyses presented in Fig 2 and S2 Fig.

### Protein Novelty Analysis

Proteins in the apicoplast proteome were manually categorized for having a potentially novel function based on PlasmoDB version 33 (released June 30, 2017) gene product annotations.

Gene products with annotations that could clearly assign a given protein to an established cellular pathway were labeled as “Known Pathway;” gene products with a descriptive annotation that did not clearly suggest a cellular pathway were labeled as “Annotated Gene Product,

Unknown Function;” and gene products that explicitly contained the words “unknown function” were labeled as “Unknown Function.”

### OrthoMCL Orthology Analysis

To analyze the conservation of candidate apicoplast proteins identified by apicoplast BioID, OrthoMCL ortholog group IDs were obtained from PlasmoDB. Based on OrthoMCL version 5 (released July 23, 2015), each ortholog group was then categorized as being present only in *Plasmodium* spp., only in Apicomplexa, or present in at least one organism outside of the Apicomplexa.

### Gene Essentiality Analysis

Genome-scale essentiality data for *P. berghei* or *P. falciparum* genes were accessed from the original manuscripts [41, 42].

### Selection of Candidates for Experimental Localization

To facilitate molecular cloning, proteins identified by BioID or PlastNN were candidates for GFP tagging only if their corresponding gene sizes were less than 2 kb. With the exception of ROM7, which was selected based on the biological interest of rhomboid proteases, we focused on localizing conserved *Plasmodium* genes of unknown function due to interest in functional characterization of the *Plasmodium* genome. PF3D7_1472800, PF3D7_0521400, and PF3D7_0721100 (all from the BioID list) were chosen due to their diverse apicoplast:ER enrichment rankings in the BioID list (S1 Table; PF3D7_1472800 ranked number 52/187 and was identified exclusively in BioID-Ap samples; PF3D7_0521400 ranked number 131/187 and was found in both samples but was enriched nearly 400-fold in BioID-Ap samples; and PF3D7_0721100 ranked 184/187 and was enriched in BioID-Ap samples only slightly above our 5-fold cutoff). From the PlastNN list, PF3D7_1349900 and PF3D7_1330100 were selected solely based on being proteins of unknown function with small gene sizes. Because of the small sample sizes of proteins selected for GFP-tagging and the non-random nature of selecting candidates, we note that the results of our experimental validation should not be extrapolated to be representative of the PPVs of the BioID and PlastNN datasets as a whole.

### Parasite Growth Time Courses

Sorbitol-synchronized ABCB7 and ABCF1 TetR-DOZI parasites were washed multiple times to remove residual ATc and were returned to culture medium containing 500 nM ATc, 200 μM IPP (Isoprenoids, LLC), or no supplements. Samples for growth assays, DNA isolation, or western blotting were harvested every other day when the majority of parasites were trophozoites and schizonts. For growth assays, parasites were fixed in 1% paraformaldehyde in PBS and were stored at 4°C until completion of the time course. Samples were then stained with 50 nM YOYO-1 and parasitemia was analyzed on a BD Accuri C6 flow cytometer. Samples for DNA isolation and western blotting were treated with 0.1% saponin in PBS to release parasites from the erythrocyte, washed in PBS, and stored at −80°C until analysis.

### Quantitative PCR

Total parasite DNA was isolated from time course samples using the DNeasy Blood & Tissue Kit (Qiagen). Quantitative PCR was performed using Power SYBR Green PCR Master Mix (Thermo Fisher) with primers CHT1 F and CHT1 R targeting the nuclear gene chitinase or TufA F and TufA R targeting the apicoplast gene elongation factor Tu (0.15 μM final concentration each primer) [46]. Quantitative PCR was performed on an Applied Biosystems 7900HT RealTime PCR System with the following thermocycling conditions: Initial denaturation 95°C/10 minutes; 35 cycles of 95°C/1 minute, 56°C/1 minute; 65°C/1 minute; final extension 65°C/10 minutes. Relative quantification of each target was performed using the ΔΔCt method.

### Statistics

95% confidence intervals were determined using the GraphPad QuickCalc for confidence interval of a proportion via the modified Wald method (https://www.graphpad.com/quickcalcs/confInterval1/). Two-way ANOVAs were performed in GraphPad Prism version 7.04.

## Acknowledgements

We thank Jacquin Niles for providing the NF54^Cas9^+^T7 Polymerase^ cell line and pSN054-V5 plasmid, Sean Prigge for α-*Pf*ACP antibody, Sebastian Mikolajczak and Stefan Kappe for α-*Pf*BiP antibody, and Walid Houry for α-*Pf*ClpP antibody. We also thank Julian Lutze for assistance with molecular cloning of candidate apicoplast genes.

## Author Contributions

Conceptualization, M.J.B. and E.Y.; Software, A.L., S.J.W., A.J., S.Z., X.W., J.Z. and S.A.R.; Investigation, M.J.B., and S.G.; Resources, L.Z., J.E.E., and S.A.R.; Writing – Original Draft, M.J.B. and E.Y.; Writing – Review & Editing, M.J.B., S.G., L.Z., A.L., S.W.J., A.J., S.Z., X.W., S.A.R., J.Z., J.E.E., and E.Y.; Supervision, E.Y., J.E.E., and J.Z.; Funding Acquisition, E.Y., J.E.E., J.Z., and S.A.R.

---

**S1 Fig.**
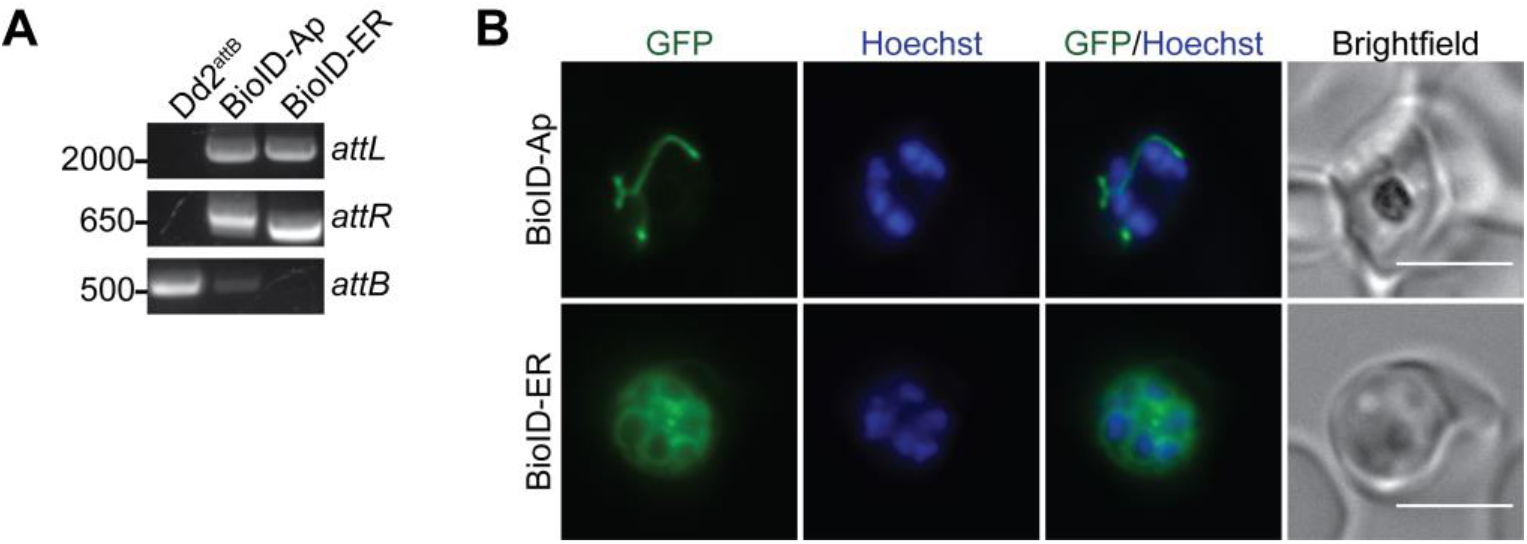
Integration and expression of BioID-Ap and BioID-ER constructs in Dd2^attB^ parasites. (A) PCR products showing integrated *attL* and *attR* sites or unintegrated *attB* site. (B) Live-cell imaging of Hoechst-stained BioID-Ap and BioID-ER parasites. Scale bars, 5 μm.

**S2 Fig.**
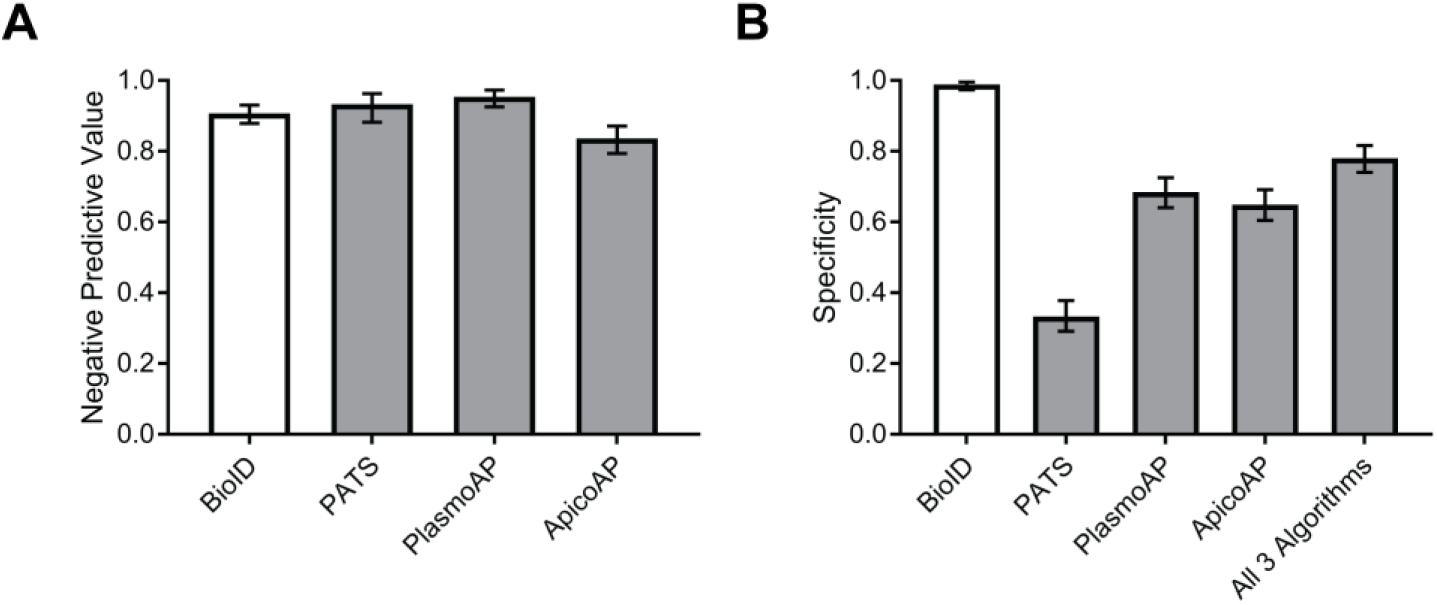
Comparison of negative predictive values and specificities of apicoplast BioID and bioinformatic prediction algorithms. (A) Negative predictive value (NPV) and (B) specificity of apicoplast BioID, PATS, PlasmoAP, and ApicoAP. Error bars represent 95% confidence intervals.

**S3 Fig.**
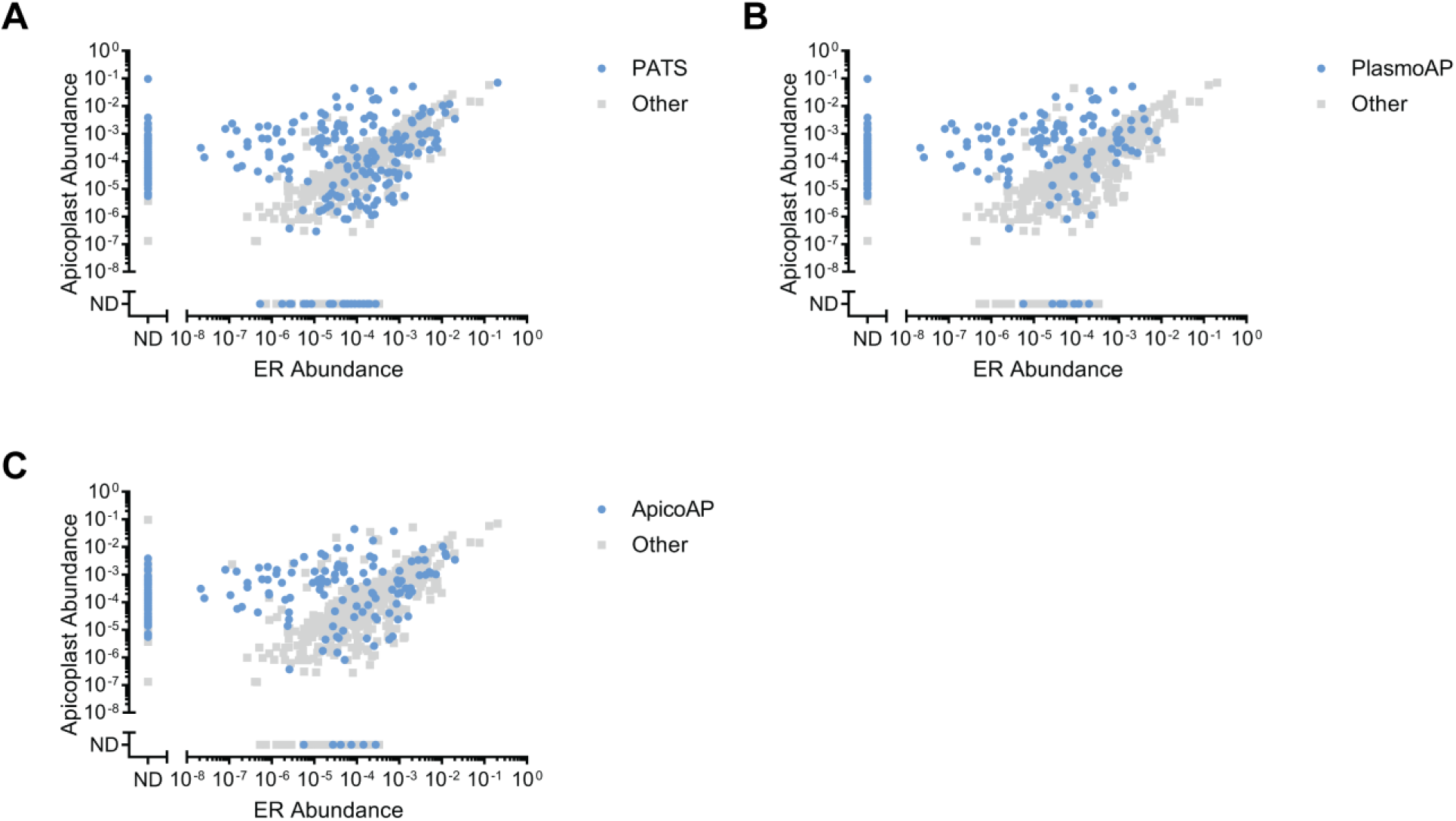
Bioinformatically predicted apicoplast proteins are not clearly distinguishable based on apicoplast:ER abundance ratio. Proteins predicted to localize to the apicoplast by (A) PATS, (B) PlasmoAP, or (C) ApicoAP are highlighted in each graph. Data points are identical to those in Fig 2A.

**S4 Fig.**
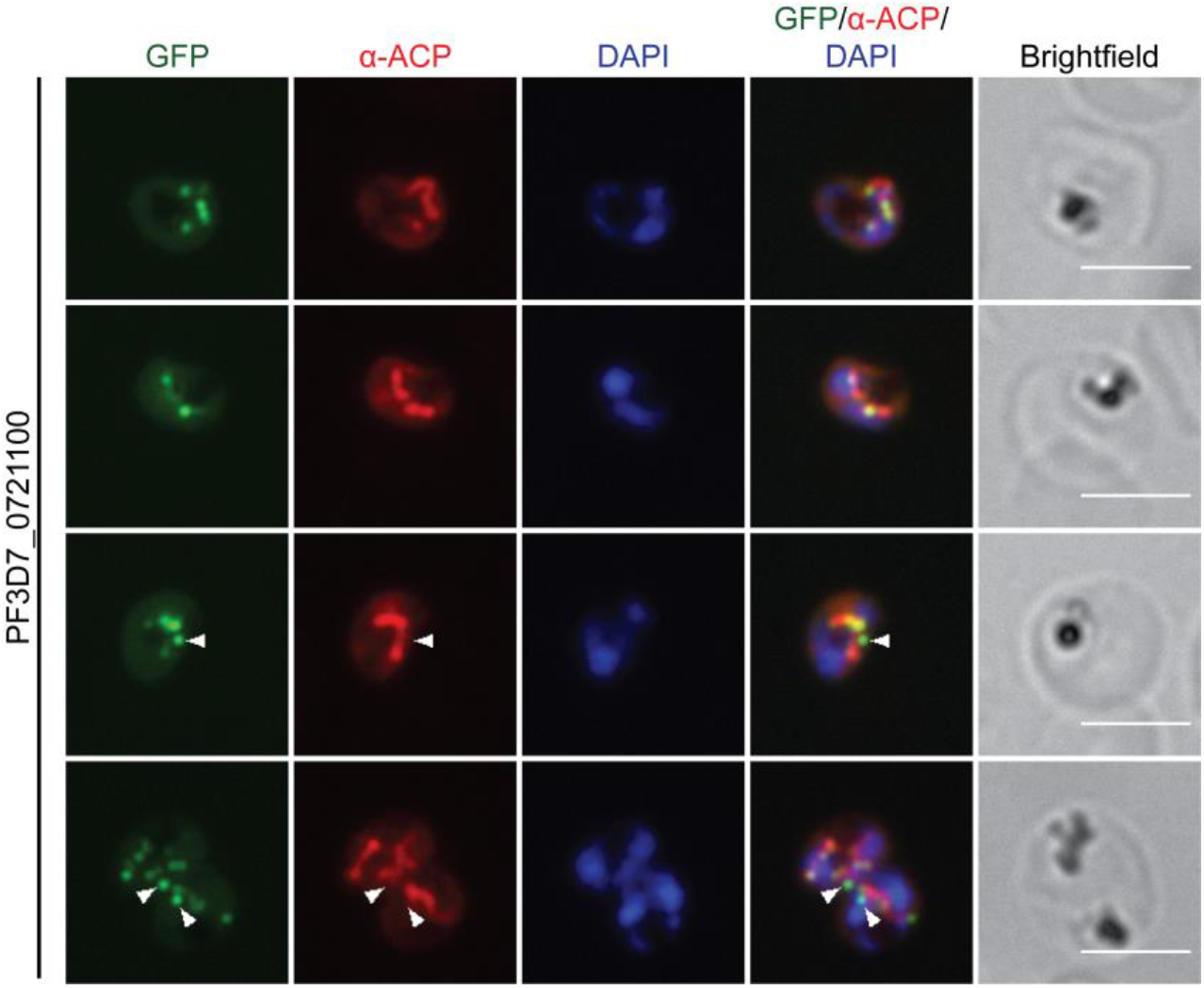
Additional fixed-cell images of PF3D7_0721100-GFP localization. PF3D7_0721100-GFP parasites were stained with an antibody against the apicoplast marker ACP. Arrowheads indicate regions where PF3D7_0721100-GFP puncta appear adjacent to as opposed to colocalizing with ACP. Scale bars, 5 μm.

**S5 Fig.**
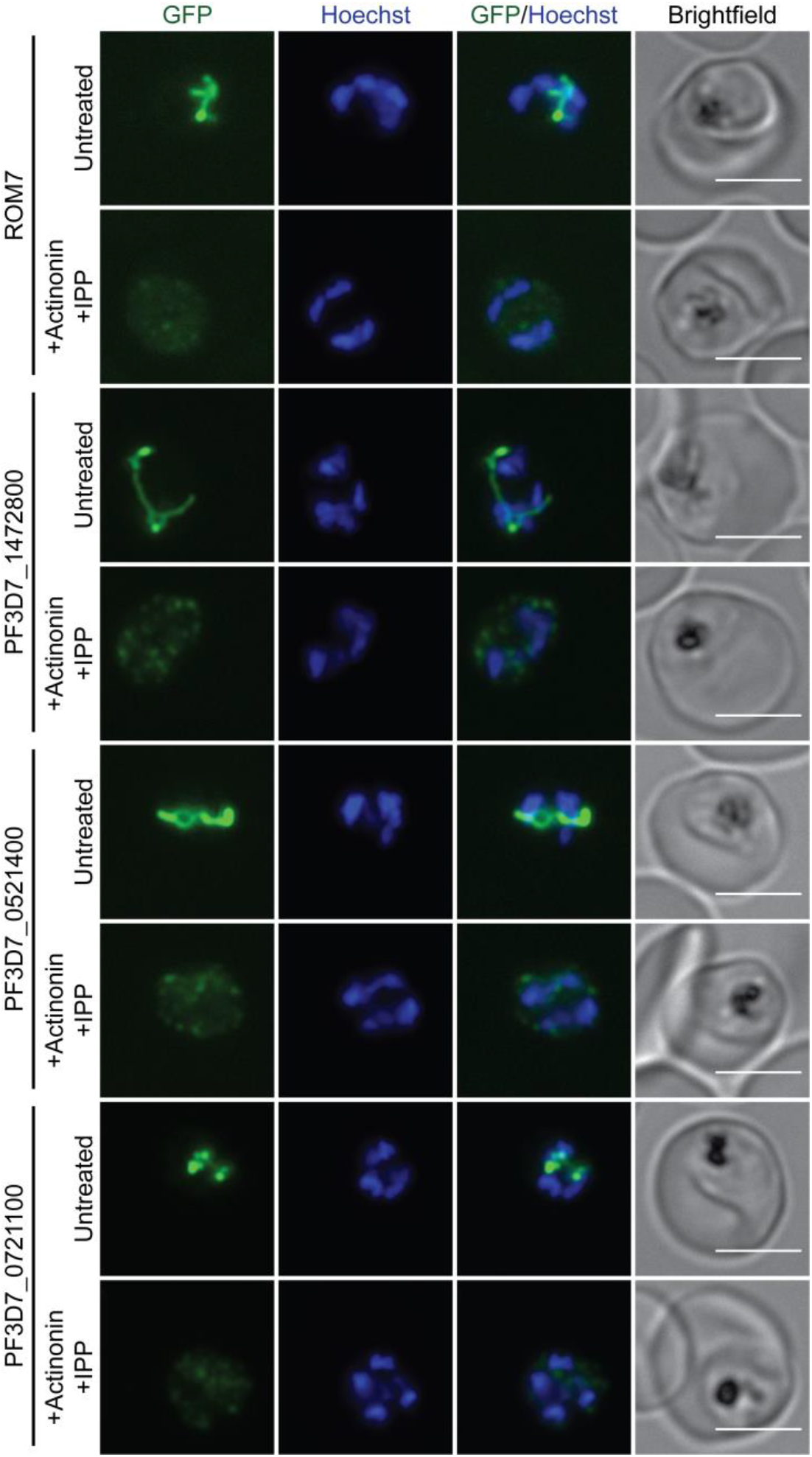
Live-cell imaging of candidate apicoplast proteins identified by BioID. Parasites expressing C-terminally GFP-tagged candidate proteins from apicoplast BioID were either untreated (apicoplast-intact) or treated with 10 μM actinonin/200 μM IPP (apicoplast-disrupted) for 3 days prior to imaging. Scale bars, 5 μm.

**S6 Fig.**
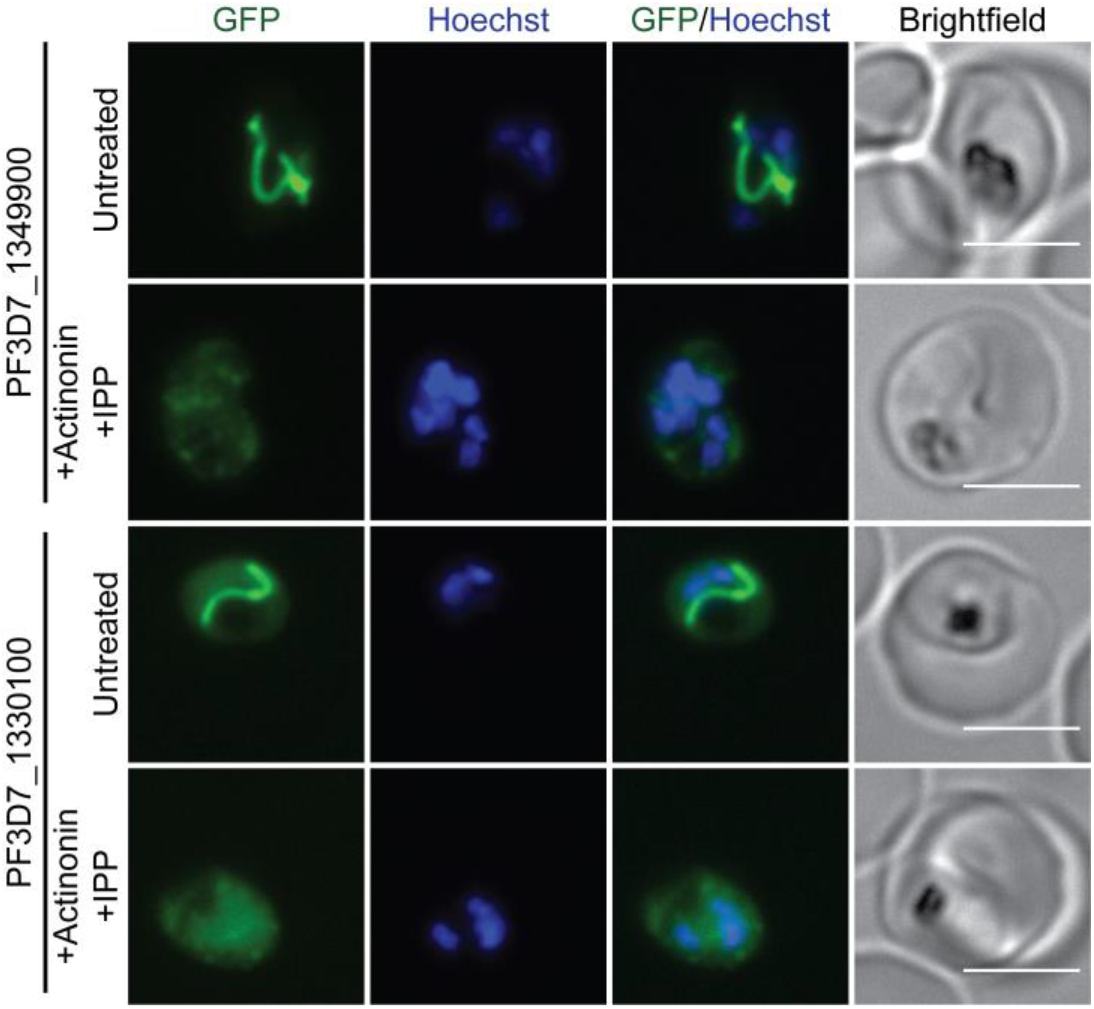
Live-cell imaging of candidate apicoplast proteins identified by PlastNN. Parasites expressing *C*-terminally GFP-tagged candidate proteins from PlastNN were either untreated (apicoplast-intact) or treated with 10 μM actinonin/200 μM IPP (apicoplast-disrupted) for 3 days prior to imaging. Scale bars, 5 μm.

**S7 Fig.**
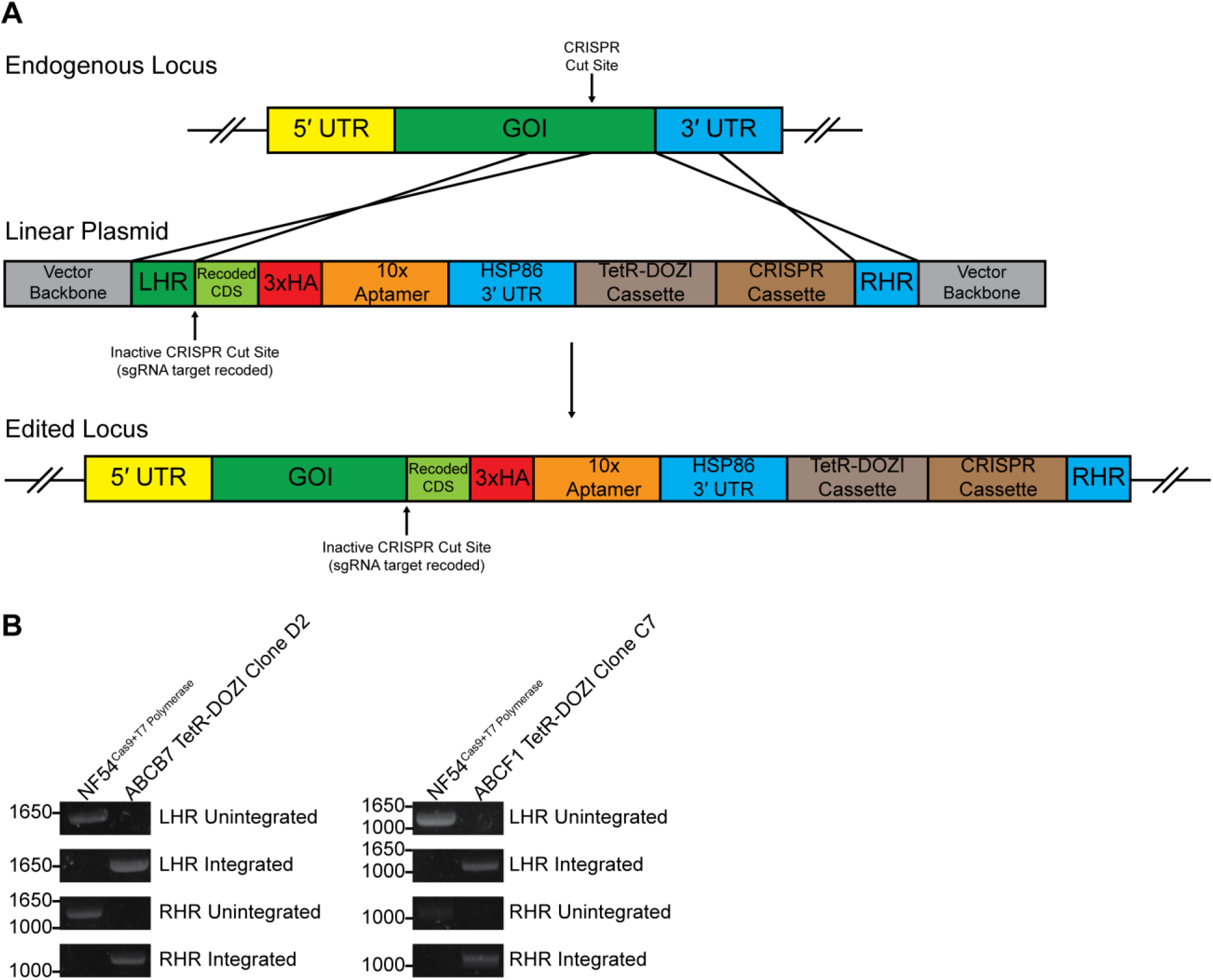
Generation of ABCB7 and ABCF1 TetR-DOZI conditional knockdown cell lines. (A) Schematic of CRISPR-Cas9-based endogenous editing to generate conditional knockdown cell lines. GOI, gene of interest; LHR, left homology region; RHR, right homology region. (B) PCR products showing integrated or unintegrated LHR and RHR sites in parental NF54^Cas9+T7 Polymerase^ or clonal genome-edited parasites.

**S8 Fig.**
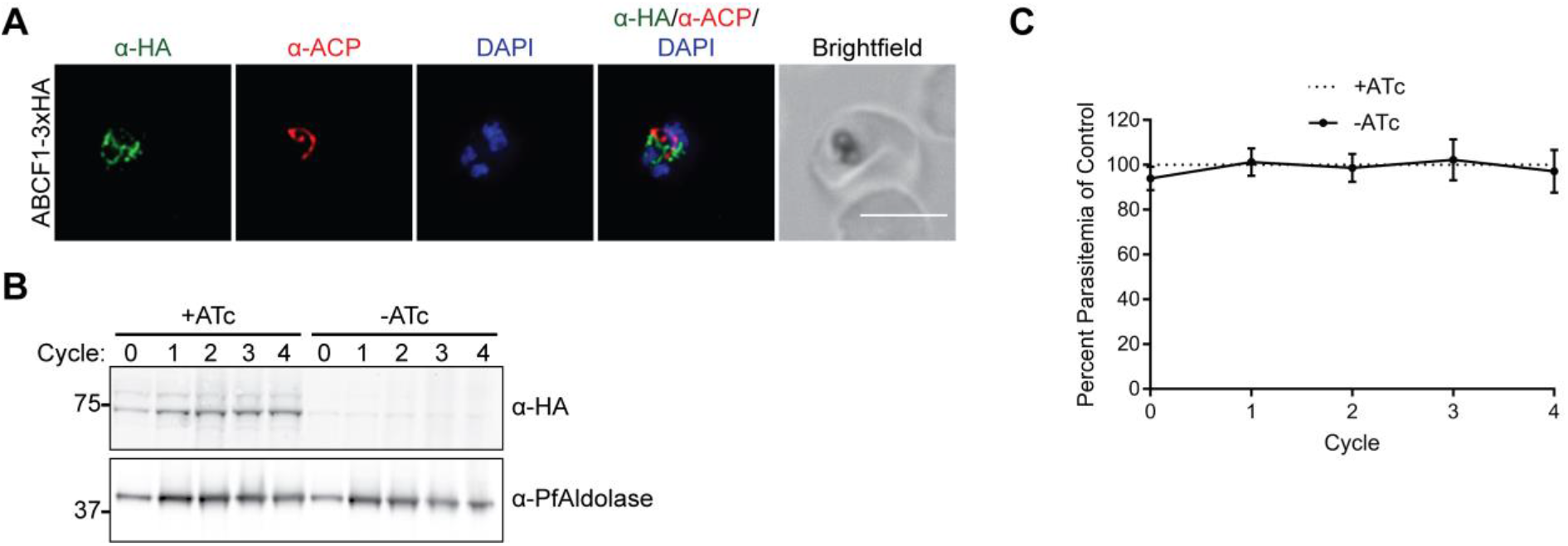
ABCB7 is a probable non-apicoplast protein for which knockdown does not cause growth inhibition. (A) Fixed-cell imaging of ABCB7-3xHA knockdown parasites stained with antibodies raised against the HA tag and the apicoplast marker ACP. Scale bar, 5 μm. (B-C) ABCB7-3xHA knockdown parasites were grown in the presence of ATc (+ATc) or the absence of ATc (-ATc) for 4 growth cycles. (B) Western blot of ABCB7-3xHA expression. (C) Parasite growth. At each time point, data are normalized to the untreated (+ATc) control. Error bars represent standard deviation from the mean of two biological replicates.

**S1 Table.** Abundances of 728 *P. falciparum* proteins identified by mass spectrometry in ≥2 biological replicates and with ≥2 unique peptides in at least one mass spectrometry run.

**S2 Table.** Positive and negative control apicoplast proteins used in this study.

**S3 Table.** Proteins predicted to localize to the apicoplast by PATS, PlasmoAP, and ApicoAP.

**S4 Table.** Positive training set used to develop PlastNN.

**S5 Table.** Layer dimensions for PlastNN neural network.

**S6 Table.** Performance of different models in cross-validation.

**S7 Table.** Results of PlastNN prediction algorithm.

**S8 Table.** Compiled list of 346 candidate apicoplast proteins based on localization in the published literature, BioID, and PlastNN.

**S9 Table.** Summary of BioID and PlastNN candidate localization data from this study and Sayers et al.

**S10 Table.** Primer and gBlock sequences used in this study.

**S1 Data.** Spreadsheet containing numerical data for Figs 2A, 2B, 2C, 2D, 3A, 3B, 3C, 5A, 5B, 5C, 5D, 8C, 8D, S2A, S2B, S7C.

